# Directed Evolution of Genetically Encoded LYTACs for Cell-Mediated Delivery

**DOI:** 10.1101/2023.11.14.567117

**Authors:** Jonathan Lee Yang, Sean A. Yamada-Hunter, Louai Labanieh, Elena Sotillo, Joleen S. Cheah, David S. Roberts, Crystal L. Mackall, Alice Y. Ting, Carolyn R. Bertozzi

**Author notes:** Corresponding authors: Carolyn R. Bertozzi, Alice Y. Ting, **Email:**. **Author Contributions:** J.L.Y., A.Y.T, and C.R.B., designed research; J.L.Y, S.Y.H, L.L, J.S.C, and D.S.R. performed research; S.Y. H, L.L, E.S. and C. L. M contributed new reagents/analytic tools; J. L. Y, S. Y. H, L.L, D. S. R, A. Y. T, and C. R. B analyzed data; and J. L. Y, A. Y. T., and C.R.B. wrote the paper. S. Y. H and L.L contributed equally. **Competing Interest Statement:** C.R.B. is a co-founder and scientific advisory board member of Lycia Therapeutics, Palleon Pharmaceuticals, Enable Bioscience, Redwood Biosciences (a subsidiary of Catalent), OliLux Bio, InterVenn Bio, GanNA Bio, Firefly Bio, Neuravid and Valora Therapeutics. S.A.Y.-H. is a consultant for Quince Therapeutics. L.L. is a cofounder of, consults for, and holds equity in CARGO Therapeutics. E.S. is a consultant for Lepton Pharmaceuticals and Galaria, and holds equity in Lyell Immunopharma. C.L.M. is a cofounder of Lyell Immunopharma, CARGO Therapeutics, and Link Cell Therapies, which are developing CAR-based therapies, and consults for CARGO, Link Immatics, Ensoma and Red Tree Capital. A.Y.T. is a scientific advisor to Third Rock Ventures and Nereid Therapeutics. The remaining authors declare no competing interests.

## Abstract

Lysosome-targeting chimeras (LYTACs) are a promising therapeutic modality to drive the degradation of extracellular proteins. However, early versions of LYTAC contain synthetic glycopeptides that cannot be genetically encoded. Here we present our designs for a fully genetically encodable LYTAC (GELYTAC), making our tool compatible with integration into therapeutic cells for targeted delivery at diseased sites. To achieve this, we replaced the glycopeptide portion of LYTACs with the protein insulin like growth factor 2 (IGF2). After showing initial efficacy with wild type IGF2, we increased the potency of GELYTAC using directed evolution. Subsequently, we demonstrated that our engineered GELYTAC construct not only secretes from HEK293T cells but also from human primary T-cells to drive the uptake of various targets into receiver cells. Immune cells engineered to secrete GELYTAC thus represent a promising avenue for spatially-selective targeted protein degradation.

**Significance Statement:** Better therapeutic windows can be achieved by targeting therapeutics to their desired sites of action. For protein therapeutics, this might be achieved by engineering cell therapies that home to a tissue of interest and secrete the biologic drug locally. Here, we demonstrate that human primary T cells can be engineered to produce genetically encoded lysosome targeting chimeras (GELYTACs). These GELYTACs mediate the degradation of extracellular proteins associated with cancer progression. Thus, cells engineered to produce GELYTACs represent a potential new class of cancer therapeutics.

## Introduction

Lysosome-targeted degradation is an emerging therapeutic modality that facilitates the degradation of membrane and soluble extracellular proteins. Compared to traditional therapeutic modalities, such as small molecule or antibody-based inhibitors, targeted protein degradation offers increased potential potency and broadens the druggable proteome (1). The first generation of this technology came in the form of lysosome-targeting chimeras (LYTACs), which are bifunctional molecules comprised of an antibody that binds to a cell surface or secreted protein of interest (POI) conjugated to a ligand that binds a lysosome trafficking receptor such as the insulin growth factor 2 receptor (IGF2R, also known as CI-M6PR) (Figure 1A) (2,3,4). Since then, other technologies have been developed, such as AbTACs (5), ProTABs (6), and KineTACs (7), among others (8,9,10,11,12,13,14,15,16,17,18, 19, 20, 21, 22). These use similar bifunctional molecules to recruit POIs either to lysosome trafficking receptors or plasma membrane-associated ubiquitin ligases and have opened a new chapter in the field of targeted protein degradation.

**Figure 1.**
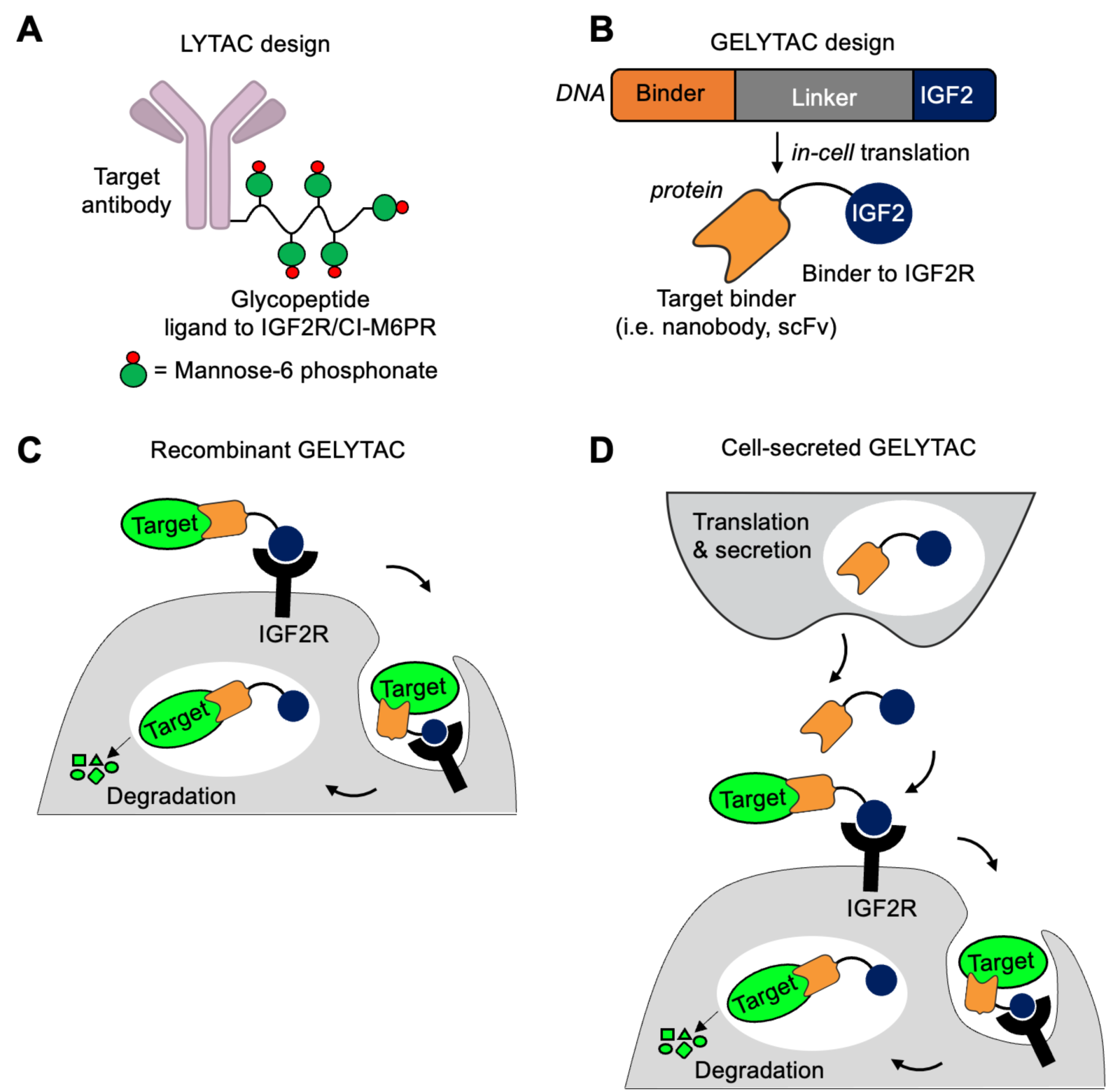
Design and applications of GELYTAC. A) Design of LYTAC (2), in which the binder to IGF2R/M6PR is a synthetic glycopeptide bearing mannose-6 phosphonate groups. LYTAC is not genetically encodable. B) Genetically Encoded LYTACs (GELYTACs) are bifunctional proteins consisting of a binder (ie nanobody or scFv) to the target of interest and IGF2, a 7.5 kDa protein that binds to IGF2R. C) GELYTACs can be utilized as a recombinant protein, where the target is internalized via GELYTAC binding to IGF2R and then degraded in the lysosome. D) Alternatively, GELYTACs can be secreted by cells (such as therapeutic T cells) to act on local targets.

There is increasing interest in new technologies that target therapeutic molecules to the sites of their desired action (23). This has recently been accomplished by encoding the production of therapeutic proteins in the genomes of targeted cell therapies. For example, therapeutic T cells or natural killer (NK) cells have been engineered to secrete scFv’s (24), bispecific T cell engagers (25, 26), cytokines (27, 28, 29), or soluble IL-12 (30) within tumor microenvironments, thereby concentrating the activity of these proteins at the desired tissue site. We are interested in developing new LYTAC constructs that can be similarly delivered.

The idea of spatially selective targeted degraders is complementary to cell-type selective LYTACs (7). However, the identification of cell-type selective lysosomal trafficking receptors remains a work in progress. Furthermore, disease states often involve multiple cell types (e.g., the tumor microenvironment which comprises a heterogeneous mixture of tumor cells, immune cells, and bystander cells). Therefore, an ideal and more effective approach should drive the degradation of the target protein by all cell types in the TME, not just a single one.

Here we report the development of Genetically Encoded LYTACs, which we term GELYTACs. These are encoded by a single transgene that can introduced into therapeutically relevant cells. The GELYTAC is comprised of two small protein modules: a small protein binder (i.e. nanobody or scFv) to a target of interest and an evolved variant of IGF2 (insulin growth factor 2) that binds to IGF2R. We improved the IGF2 scaffold via directed evolution and demonstrated its ability to selectively target extracellular mCherry, TGF-β, and shed IL6R ectodomain. To illustrate the potential utility for cell-based therapeutics, we also show that engineered primary human T cells secreting GELYTACs can effectively induce uptake of aforementioned targets by tumor cells. Our study introduces a promising new format for the effective and versatile clearing of extracellular proteins by degraders secreted by engineered cells. This could potentially be later applied for the spatially selective degradation of targets in the tumor microenvironment.

## Results

### Design of GELYTACs

To design an all-protein, genetically-encodable bifunctional molecule for targeted degradation of extracellular proteins, we fused a small protein binder via a flexible linker to IGF2 peptide (Figure 1B). IGF2 binds to domain 11 of IGF2R with a K_D_ of 4.5 nM (31) and is subsequently shuttled to the lysosome (32, 33). This biologic can either be administered recombinantly or be delivered by therapeutic cells as a cell therapy (Figure 1C-D).

As a proof of concept, we designed and generated GELYTAC targeting a model protein, mCherry. We selected the nanobody LAM4 which binds to mCherry with a K_D_ of 180 pM (34) and fused it to IGF2 (Figure 2A). We were able to easily produce mCherry GELYTAC by recombinant expression in *E. coli*. To test mCherry GELYTAC’s ability to selectively target and mediate internalization of mCherry into cells, we treated K562 leukemia cells with a mixture of mCherry protein (100 nM) and mCherry GELYTAC (from 0-1000 nM) for 24 hrs (Figure 2B). Since mCherry is more resistant to lysosomal degradation (35), we expect GELYTAC to drive fluorescent mCherry accumulation into cells’ lysosomes. Flow cytometry showed a 12-fold increase in mCherry fluorescence in K562 cells at 275 nM of mCherry GELYTAC while controls with mCherry nanobody only or an alternate-targeting GELYTAC (EGFR GELYTAC) showed no mCherry uptake in K562 cells across all concentrations. In agreement with the flow cytometry data, fluorescence microscopy showed that mCherry colocalized with stained lysosomes only in the presence of mCherry GELYTAC and not in the controls (Figure 2C, Figure S1B). A time course of mCherry uptake revealed maximum uptake after 24 hrs of mCherry GELYTAC treatment (Figure S1A).

**Figure 2.**
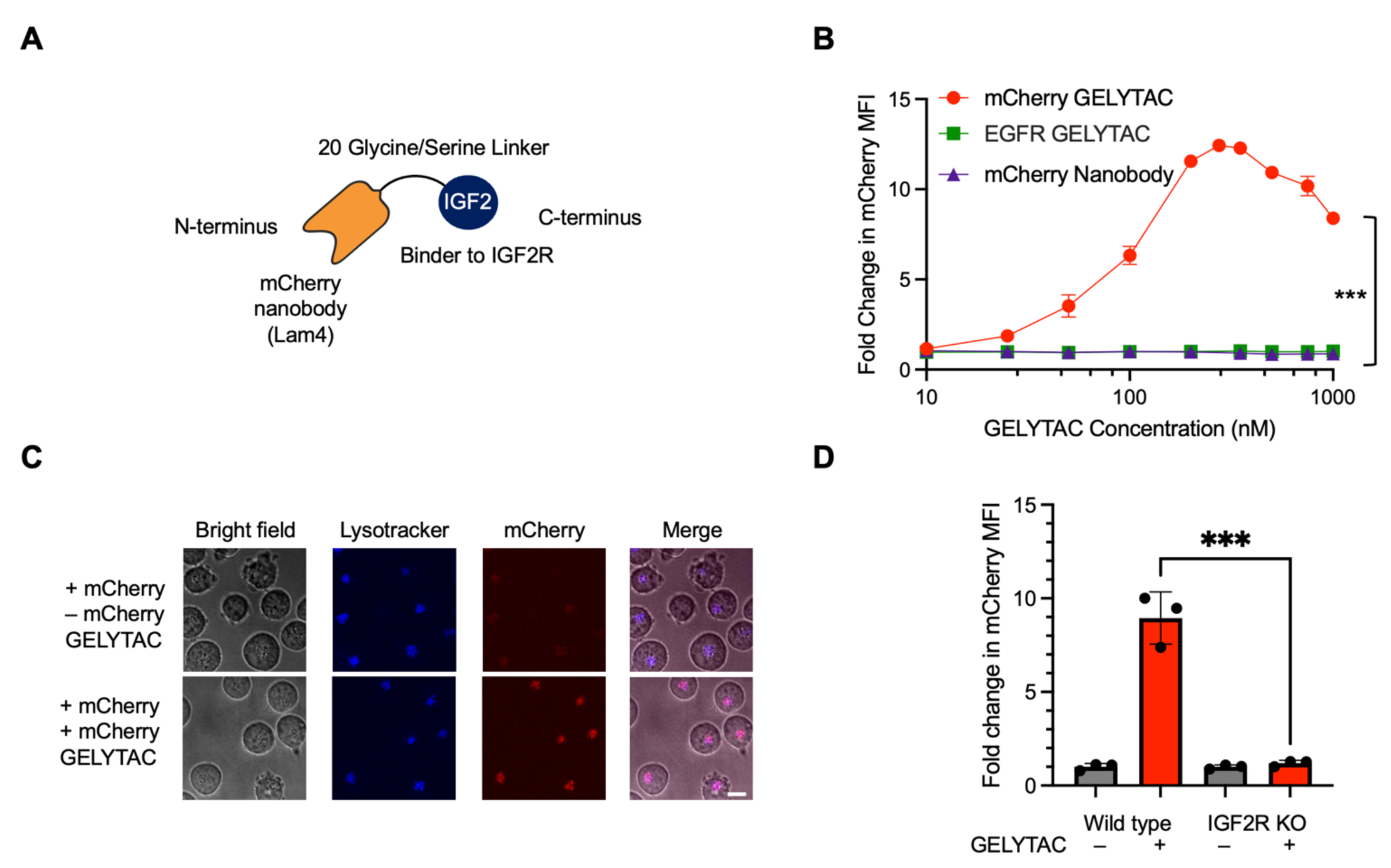
Recombinant GELYTACs mediate internalization of mCherry by K562 cells. A) Design of mCherry GELYTAC. LaM4 is a nanobody that binds to mCherry with a K_D_ of 180 pM^16^. mCherry internalization as a function of GELYTAC concentration. Median fluorescence intensity (MFI) of K562 cells was measured 24 hrs after treatment with mCherry GELYTAC (circles) and recombinant mCherry (100 nM). Negative controls are mCherry nanobody only (triangles) and a GELYTAC targeting EGFR (squares). Errors bars represent the standard deviation (SD) from 3 biological replicates. *** = p < 0.001 (determined using parametric t-test). C) Confocal fluorescence images of live K562 cells after 1-hr treatment with 100 nM mCherry and 275 nM mCherry GELYTAC. Scale bars, 10 μm. D) GELYTAC-mediated mCherry internalization by wild type and IGF2R KO K562 cells. The experiment was performed at 275 nM of GELYTACs and 100 nM mCherry for 4 hrs. Errors bars represent the SD from 3 biological replicates. *** = p < 0.001 (determined using parametric t-test).

To elucidate the mechanism of action, we repeated the flow cytometry experiments using IGF2R knockout K562 cells. In these samples, no uptake of mCherry was observed (Figure 2D) in the presence of mCherry GELYTAC. These data suggest that GELYTAC effectively targets soluble extracellular protein for lysosomal targeting of via recruitment of IGF2R.

### Directed evolution of GELYTACs

We designed GELYTAC such that they can be secreted by therapeutic cells. However, unlike recombinant proteins, for which we can specify dosing concentrations, we cannot control secreted GELYTAC concentrations. Thus, we aimed to make GELYTACs more effective at lower concentrations by lowering their EC_50_ through directed evolution, which can be streamlined due to the GELTYAC being genetically encodable.

We hypothesized that by evolving the IGF2 peptide in GELYTACs, we would be able to select for mutants with tighter binding to IGF2R and consequently, more potent POI internalization and degradation. To test whether higher affinity IGF2 mutants translates to a more efficacious GELYTAC, we first created mCherry GELYTACs using a series of known point mutants of IGF2 with varying affinities for IGF2R (Figure 3A) (31, 36). We observed a positive correlation between IGF2 variants’ K_D_’s and mCherry internalization by the corresponding mCherry GELYTAC (Figure 3B). This provided the rationale to engineer an improved IGF2 binder for improving GELYTAC potency through yeast surface display directed evolution.

**Figure 3.**
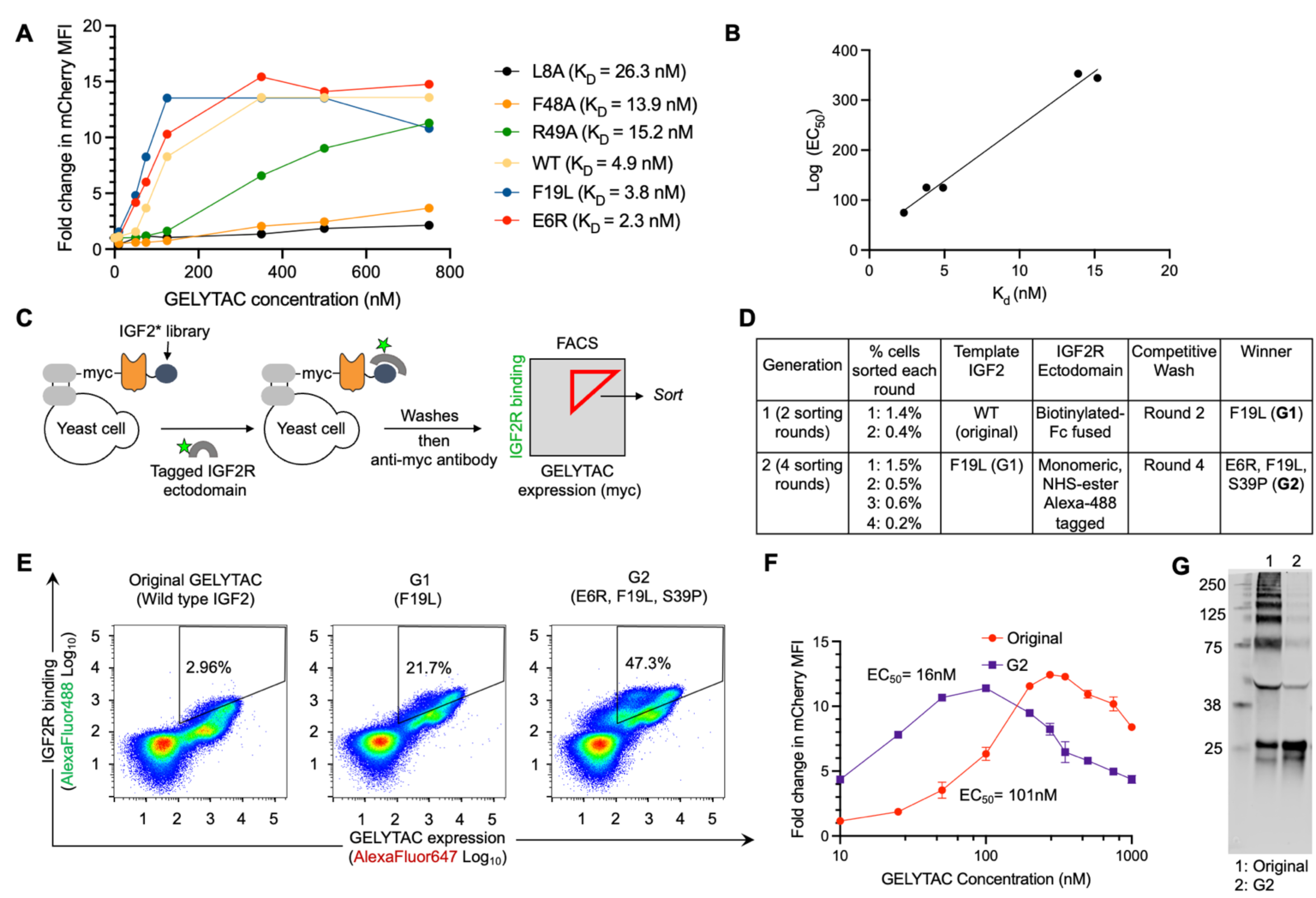
Directed evolution of GELYTAC. A) The effect of point mutations in IGF2 on GELYTAC-mediated internalization of mCherry. mCherry uptake assay was performed as in Figure 2B. B) Log(EC_50_) calculated from the data in Figure 3A plotted against published K_D_ values (32, 37). C) mCherry GELYTAC library is displayed as a fusion to Aga2p. Yeast cells were treated with AlexaFlour488-labeled IGF2R ectodomain (100 nM), washed 3 times, then stained with anti-myc antibody and sorted by FACS. D) Summary of selection conditions used across two generations and 6 rounds of sorting. E) 2D flow cytometry analysis of yeast expressing original and evolved mCherry GELYTACs (G1 and G2). Cells were stained with IGF2R ectodomain and anti-myc antibody as in Figure 3C, then washed and analyzed by flow cytometry. F) GELYTAC-mediated internalization (original and G2) of mCherry by K562 cells, performed as in Figure 2B. G) Anti-FLAG Western blot of original and G2 mCherry GELYTACs secreted from HEK293T cells. SDS-PAGE was run under non-reducing conditions to reveal oligomerization of original GELYTAC and reduced oligomerization of G2 GELYTAC.

Using the mCherry GELYTAC as a scaffold, we introduced diversity in the IGF2 domain by error prone PCR. The library was displayed on the yeast cell surface via fusion to the C-terminus of the yeast mating protein Aga2p (37, 38, 39, 40). To select for clones capable of binding to IGF2R with high affinity, we incubated the yeast library with tagged recombinant IGF2R (Fc-fused or monomeric). Staining with anti-myc antibody was used to quantify GELYTAC expression level. Two-dimensional fluorescence-activated cell sorting (FACS) sorting was used to enrich library members with high IGF2R/anti-myc intensity ratio (Figure 3C). To increase selection stringency in the last round of generation 1 and generation 2, we modulated our staining protocols to include either competitive washes or eliminate the avidity effects from Fc-fused IGF2R by shifting to monomeric IGF2R (Figure 3D). Ultimately, after 6 rounds of sorting over 2 generations of libraries, we isolated a clone with the IGF2 mutations E6R, F19L, and S39P (termed ‘G2’), exhibiting improved binding to IGF2R’s ectodomain at expression levels matched to that of wild-type IGF2 (Figure 3E, Figure S2A-C). This mutant contained two published mutations (31, 36) known to lower IGF2’s K_D_(E6R and F19L), as well as a novel mutation, S39P. We verified that S39P is a crucial mutation for G2’s improved binding by comparing G2’s triple mutant to the E6R and F19L double mutant (Figure S2E).

To verify that the benefits of directed evolution translate into a cellular context, we compared the efficacy of the wildtype GELYTAC (original GELYTACs) to our evolved G2 GELYTAC in mCherry uptake by K562 cells and saw drastic reduction of the EC_50_ from 101 nM to 16 nM (Figure 3F, Figure S3B). Finally, to see if both the original and G2 GELYTACs can be secreted by HEK293T cells, we ran an anti-FLAG western blot of the supernatant of HEK293T cells transfected with these constructs. Not only did both constructs secrete, but the evolved G2 GELYTAC also exhibited decreased propensity to oligomerize (Figure 3G, Figure S4B-C). This increases the amount of biologically active material, which is likely to contribute to improvements in the in efficacy of downstream cell-based therapy applications where GELYTACs cannot be purified.

### Cell-based secretion of GELYTACs and knockdown of other targets

Next, we explored an alternate mode of GELYTAC administration: secretion by a HEK293T sender cell to uptake soluble proteins into a receiver K562 cell (Figure 4A). To achieve this, we cocultured HEK293T cells transfected with mCherry GELYTAC (original or G2), control GELYTAC (alternate-targeting, IL6R GELYTAC), or mCherry nanobody only with GFP-expressing K562 cells for 24 hrs in media supplemented with 100 nM mCherry (Figure 4B). We verified the presence of all constructs in the conditioned media at 24 hrs by Western blot and observed comparable secretion amongst the constructs (Figure S4A). We also observed another band of slightly lower mass in both GELYTAC secretion samples. To verify the identity of the second band, we analyzed both GELYTACs by top-down mass spectrometry; while both contained a portion of truncated GELYTAC (truncation of 28 and 27 amino acids for original and G2, respectively), the predominant species corresponded to the full length (Figure S5A-D).

**Figure 4.**
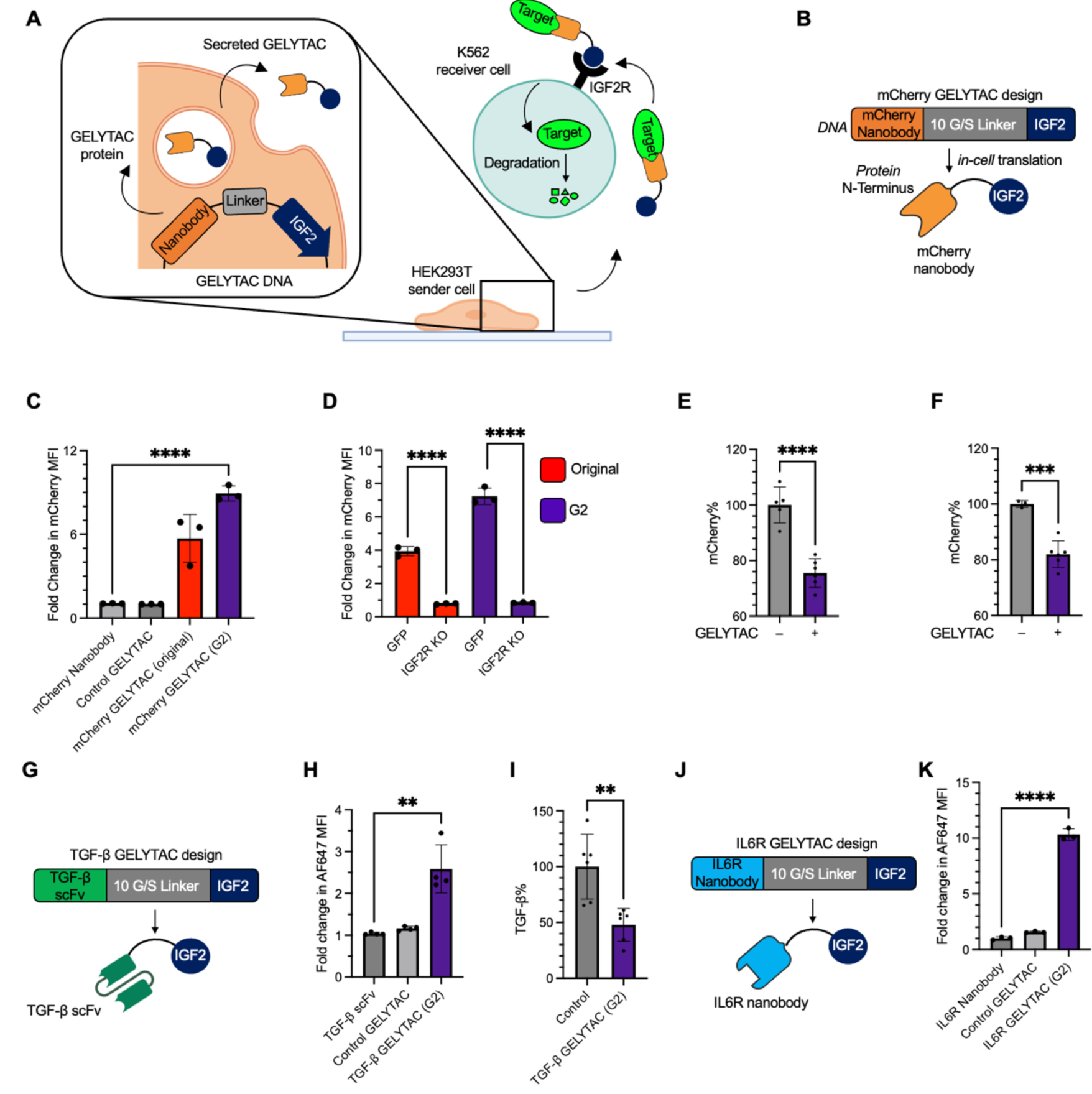
Delivery of GELYTAC via cell secretion. A) Schematic of cocultured sender and receiver cells. Adherent HEK293T cells secrete GELYTAC that acts on suspended K562 cells to internalize and degrade targets. B) Design of mCherry GELYTAC used in coculture experiments. Cocultured cells were treated with mCherry for 24 hrs. Then, K562 receiver cells were separated and analyzed by flow cytometry. HEK293T secreting either original or G2 GELYTAC were tested. Control GELYTAC targets IL6R instead of mCherry. Errors bars represent the SD from 3 biological replicates. **** = p < 0.0001 (determined using parametric t-test). D) Cocultures comprising either wild type or IGF2R KO K562 cells were treated with mCherry for 24 hrs and analyzed as in Figure 4C. E) To quantify mCherry clearance in the coculture system, media was analyzed for mCherry fluorescence 72 hrs after mCherry addition using a plate reader. This experiment was repeated 5 times. Errors bars represent the SD from 5 biological replicates. **** = p < 0.0001 (determined using parametric t-test). F) To quantify mCherry degradation, both cells and media were collected and analyzed for mCherry fluorescence after mCherry addition using a plate reader. This experiment was repeated 5 times. Errors bars represent the SD from 5 biological replicates. *** = p < 0.001 (determined using parametric t-test). G) Design of TGF-β GELYTAC utilizing a TGF-β scFv derived from the clinical candidate Fresolimumab (43). H) Cocultured cells were treated with AlexaFluor-647 tagged TGF-β for 24 hrs. Then, K562 receiver cells were separated and analyzed by flow cytometry. HEK293T cells secreting G2 GELYTAC were tested. Control GELYTAC targets mCherry instead of TGF-β. Errors bars represent the SD from 3 biological replicates. ** = p < 0.01 (determined using parametric t-test). I) To quantify TGF-β (supplemented biotinylated TGF-β) degradation, both cells and media were collected and analyzed for biotin signal via streptavidin-800 blot. This experiment was repeated 6 times. Errors bars represent the SD from 6 biological replicates. ** = p < 0.01 (determined using parametric t-test). J) Design of IL6R GELYTAC utilizing an IL6R nanobody derived from clinical candidate ALX-0061 (44). K) Cocultured cells were treated with AlexaFluor-647 tagged IL6R for 24 hrs, then K562 receiver cells were separated and analyzed by flow cytometry. HEK293T cells secreting G2 GELYTAC were tested. Control GELYTAC targets mCherry instead of IL6R. Errors bars represent the SD from 3 biological replicates. *** = p < 0.001 (determined using parametric t-test).

Next, we analyzed the receiver cells using flow cytometry and observed a ∼6-fold and ∼9-fold increase in mCherry median fluorescence intensity in K562 cells cocultured with HEK293Ts secreting original and G2 mCherry-GELYTACs, respectively (Figure 4C). We then performed the same experiment with the original and G2 mCherry GELYTACs on IGF2R KO K562 cells and saw no apparent uptake by the IGF2R KO cells, demonstrating that uptake of proteins in the coculture model is mediated by IGF2R, as we observed for treatment with the recombinant GELYTAC.

While we had observed robust mCherry uptake into receiver cells as a proxy for protein degradation, to determine if both secreted mCherry GELYTACs could mediate effective clearance and degradation of mCherry, we analyzed mCherry fluorescence in the coculture supernatant for mCherry clearance and the combined supernatant plus cells for mCherry degradation using a plate reader. After incubation of GELYTAC secreting HEK293T cells with K562 receiver cells, we observed that ∼25% of mCherry is cleared (Figure 4E) from the media and ∼20% of mCherry is degraded (Figure 4F).

Finally, we demonstrate that the GELYTAC design we optimized for targeting mCherry can be generalized towards the internalization of more therapeutically relevant targets by simply exchanging the nanobody. To this end, we developed GELYTACs targeting soluble proteins TGF-β and IL6R. TGF-β and IL6R are immunosuppressive factors in the tumor microenvironment, so depleting these soluble factors could help improve the efficacy of cancer immunotherapy treatments, such as immune-checkpoint blockade and CAR-T therapy (23). Local degradation of these targets is attractive because systemic targeting of immunosuppressive factors has been shown to be toxic (41). We designed the TGF-β GELYTAC by fusing our engineered IGF2 with a TGF-β scFv derived from the clinical candidate antibody Fresolimumab (42) (Figure 4G). We tested the TGF-β GELYTAC in a coculture system with HEK293T secreting the TGF-β GELYTAC and GFP-expressing K562 and supplemented the media with AlexaFluor647 (AF-647)-tagged TGF-β. Like the mCherry assays, we observed ∼2.6-fold increase in AF-647 fluorescence in K562’s (Figure 4H, Figure S4B). We also analyzed if TGF-β GELYTAC could mediate the degradation of TGF-β. To do this, we added 1 nM of biotinylated TGF-β to cocultured K562 cells with HEK293T cells secreting either mCherry GELYTAC and TGF-β scFv or TGF-β GELYTAC and analyzed via a streptavidin western blot of a mixture of the cell lysate and supernatant. We observed that the TGF-β GELYTAC mediated ∼52% reduction in TGF-β when compared to the combined mCherry GELYTAC and TGF-β scFv control (Figure 4I, Figure S4C). For IL6R, we developed a GELYTAC containing engineered IGF2 fused to a clinical candidate IL-6R nanobody (ALX-0061) (43) (Figure 4J). In the coculture assay, we supplemented the media with AF-647-tagged IL-6R and saw ∼10-fold increase in AF-647 signal in the K562 cells when cocultured with IL6R GELYTAC secreting HEK293T cells (Figure 4K, Figure S4D). In these experiments, the MFI increase mediated by secreted TGF-β GELYTAC was lower compared to mCherry and IL6R GELYTACs. This could be due to binding of TGF-β to the TGF-β1 receptor or other receptors resulting in increased background signal. Development of tighter binding nanobodies could thus improve efficacy of this class of GELYTAC that compete with cell-surface receptors for target binding. Overall, these results demonstrate the modularity and potential of GELYTAC to recognize a wide range of targets by simply exchanging the POI binder.

To potentially enable spatial specificity for GELYTACs, we imagined that GELYTAC can be integrated into adoptively transferred T cell therapy, such as CAR-T therapy. In adoptive T cell therapy, engineered T cells home to tumors and proliferate in the tumor microenvironment based on recognition of specific tumor antigens. Furthermore, T cells can be engineered to secrete factors under the control of logic gates determined by recognition of tumor-specific factors. Thus, secretion from T cells could enable spatial specificity at the tumor site. To demonstrate the feasibility of this approach, we interrogated the potential for primary human T cells to secrete GELYTACs (Figure 5A). To do this, we transduced donor human primary T cells with retrovirus encoding for mCherry G2 GELYTAC. We observed robust GELYTAC secretion (Figure S6A), and upon co-incubation of GELYTAC secreting T cells with K562 cells, observed 5.25-fold and 2.2-fold increases in mCherry MFI in K562 and T cells, respectively (Figure 5B-C). These results demonstrate the potential for secreted GELYTAC robustly on multiple cell types within an environment. Additionally, the ability of sender cells to act on themselves opens the possibility for feedback to regulate their GELYTAC secretion.

**Figure 5.**
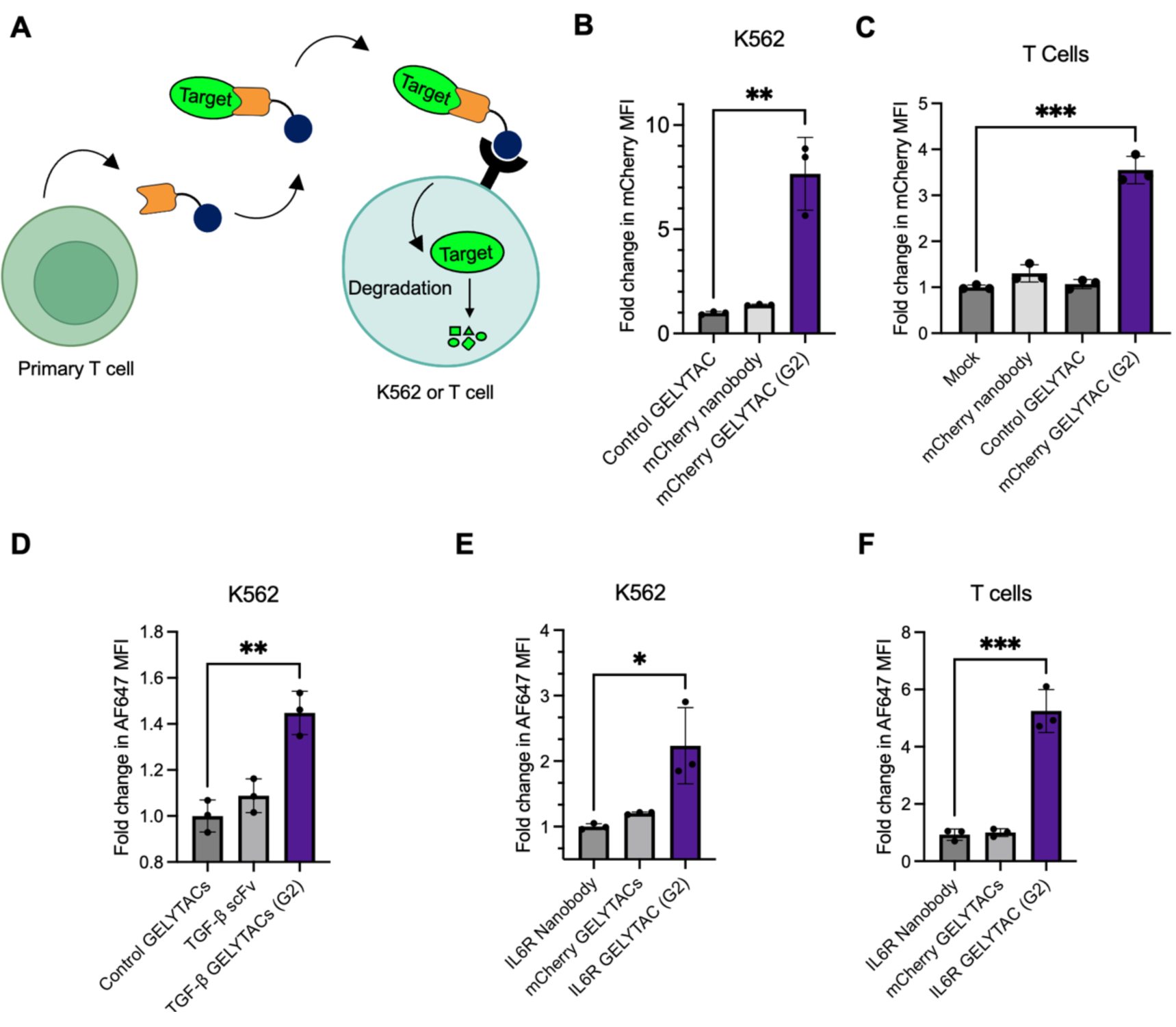
GELYTACs secreted from primary T cells mediate uptake of multiple targets. A) Schematic of T cell coculture system. Adherent human primary T cells secrete GELYTAC that acts on suspended K562 cells and T cells to internalize and degrade targets. B) Cocultured cells were treated with mCherry for 62 hrs. Then, K562 cells were analyzed by flow cytometry. T cells secreting G2 GELYTAC were tested. Control GELYTAC targets IL6R instead of mCherry. Errors bars represent the SD from 3 biological replicates. *** = p < 0.001 (determined using parametric t-test). C) Experiment performed as in (B) but measuring fold change in mCherry MFI in T cells from the coculture system. Errors bars represent the SD from 3 biological replicates. *** = p < 0.001 (determined using parametric t-test). D) Cocultured cells were treated with AlexaFluor-647 tagged TGF-β for 52 hrs. Then, K562 receiver cells were analyzed by flow cytometry. T cells secreting G2 GELYTAC were tested. Control GELYTAC targets mCherry instead of TGF-β. Errors bars represent the SD from 3 biological replicates. ** = p < 0.01 (determined using parametric t-test). E) Cocultured cells were treated with AlexaFluor-647 tagged IL6R for 9 hrs. Then, K562 receiver cells were analyzed by flow cytometry. T cells secreting G2 GELYTAC were tested. Control GELYTAC targets mCherry instead of IL6R. Errors bars represent the SD from 3 biological replicates. * = p < 0.1 (determined using parametric t-test). F) Experiment performed as detailed in (E) but measuring fold change in AlexaFluor-647 MFI in T cells from the coculture system. Errors bars represent the SD from 3 biological replicates. *** = p < 0.001 (determined using parametric t-test).

Lastly, we tested TGF-β and IL6R GELYTACs in the primary T cell coculture system. For TGF-β GELYTACs, we observed a 1.6-fold increase in TGF-β uptake into K562 cells (Figure 5D, Figure S6B). For IL6R GELYTACs, we observed 2.2-fold increase and 5.2-fold increase in IL6R uptake in K562s and T cells, respectively (Figure 5E–F, Figure S6C).

## Discussion

In this study, we developed a small, genetically encoded LYTAC that can be secreted by primary human T cells. While there are multiple approaches to make a genetically encoded LYTAC, a major benefit of using IGF2 is that it is a human protein, which should decrease its immunogenicity compared to nonhuman mammalian proteins or computationally (44,45) designed proteins (46). Indeed, during our study, another paper was published that utilized a protein-based targeting chimera composed of 2 computationally designed binders to internalize EGFR via cell secretion (21), which could be immunogenic. Other work was also published during the preparation of this manuscript that utilizes IGF2-binder fusions but does not contain cell therapy applications or directed evolution (22).

To improve the potency of GELYTACs as mediators of extracellular protein degradation, we evolved a mutant IGF2 that binds more strongly to IGF2R. Using directed evolution, we derived an IGF2 variant that, when integrated into a GELYTAC, was approximately 10-fold more potent in mediating mCherry internalization. An added benefit of the improved GELYTAC was its higher expression levels in the HEK293T/K562 coculture model and its lower propensity to aggregate. This suggests that the improved performance of the evolved GELYTAC not only stems from increased potency as a biologic but also from increased levels of active species.

We engineered primary T cells to secrete GELYTACs as a model for a future CAR-T therapy. CAR-T cells are designed to home to and proliferate within tumors and have been shown to deliver protein cargos efficiently to tumors *in vivo* (25, 27). Thus, we believe that CAR-T cells will be capable of delivering GELYTACs to the tumor microenvironment. The GELYTACs we report here have low molecular weights (20-30 kDa), and therefore likely short serum residence times (47). Therefore, we would expect GELYTACs that have diffused away from the targeted environment to be rapidly cleared from circulation. The concept of spatial selectivity by small biologics secreted by T cells has been demonstrated using bi-specific T cell engagers (24).

Finally, the GELYTAC platform synergizes with several recent technologies for therapeutic development. For example, computational methods for de novo protein design as illustrated by Baker and coworkers’ recent work (21) can be deployed to optimize GELYTACs or to add additional functionality. Lastly, because they are genetically encoded, GELYTACs can be delivered via other forms of genetic medicine such as mRNA and viral gene therapy vectors.

## Materials and Methods

### Recombinant GELYTACs production

BL21 DE3 *E. coli* (Agilent) were transformed with a vector containing sequences encoding for GELYTACs or nanobody only controls with pelB signal sequence for localization to the periplasm for disulfide bond formation. A colony was picked into 10mL of LB (supplemented with 2% glucose and kanamycin) overnight at 37 °C. The next day, the starter culture was added to 1L of 2xYT (supplemented with antibiotics) and grown at 37 °C to an OD600 ∼ 1.0-1.3. The culture was then induced at 1mM and grown overnight at 16 °C at 225 rpm. The next day, the culture was centrifuged at 7000g for 10 min and the supernatant discarded. The cell pellet was then resuspended thoroughly with 20mL 1x TES (0.2M Tris, pH=8, 0.5 mM EDTA, 0.5 M sucrose) and then the mixture was added to 20mL of ice cold ddH_2_O (supplemented with protease inhibitor). The mixture was then incubated overnight at 4 °C with shaking. The next day the mixture was centrifuged at 16,000xg. The supernatant was then filtered using a 5uM filter and then purified using Ni NTA column (Cytiva/ GE Healthcare) on an FPLC (AKTA Pro). Following Ni-NTA purification, the mixture was then purified using size exclusion chromatography (Superdex 75 Increase 10/300 GL) to and only the monomer was isolated.

### Yeast display

The yeast culture and display protocols and library generation protocols are described in depth in this publication ^35^ and in the supplemental information.

### Cell Culture (excluding primary T cell culture)

All cell lines used were less than passage 20. HEK 293T cells (ATCC) and 293GP retroviral packaging line (gift from Surgery Branch (National Cancer Institute, National Institutes of Health)) were cultured in a DMEM (Gibco) supplemented with 10% fetal bovine serum (FBS), 1% Glutamax (Gibco), 100 units/mL penicillin, and 100 mg/mL streptomycin at 37 °C under 5% CO2. K562’s (ATCC) was cultured in in RPMI (Sigma Aldrich) supplemented with 10% FBS, 1% Glutamax (Gibco), 100 units/mL penicillin, and 100 mg/mL streptomycin at 37 °C under 5% CO2.

### HEK293T coculture with K562

12 well plates were coated with HFN (1mL of HFN from Sigma added to 50mL PBS), by adding 700uL of PBS to the wells. A 70-90% confluent HEK293T T75 flask were lifted, by aspirating media and adding ∼1-2mL trypsin. Cells were diluted to 3.5e5 cells/ mL, and 1mL was added to each well and slightly agitated before returning it in the incubator. On the next day, cells were transfected with plasmids encoding for GELYTACs or controls.

To make the transfection mix, 1000ng of plasmids encoding for GELYTACs or controls were added to 100uL of blank DMEM. The mixture was mixed by flicking the tube, and then 5uL of polyethylene imine (PEI). The tube was flicked gently to homogenize, and allowed to incubate for 20 min. After incubation, the transfection mix was added directly to cells dropwise, cells were returned to incubator for 12-18 hrs. After 12-18 hrs, media + the transfection mix were aspirated and replated with 1 mL of RPMI mCherry with 3.0e5 GFP K562/mL. After 3 hrs 100nM of soluble antigen (ie mCherry, TGF-β, or IL6R) was spiked into the coculture.

At the time of analysis, the GFP K562’s was analyzed by flow cytometry for median fluorescence of mCherry or AlexaFluor-647 tagged proteins. GFP was used to distinguish between K562 cells and HEK293T cells.

### Retrovirus production

Retroviral supernatant was packaged using 293GP cells and the RD114 envelope plasmid. In brief, 11μg RD114 and 22μg of the corresponding MSGV1 transfer plasmid that contain GELYTACs or controls were delivered to 293GP cells grown on 150mm HFN dishes (Corning) to 80% confluency by transient transfection with Lipofectamine 2000 (Thermo Fisher). Media was replenished every 24 hrs. <u>Virus</u> production was performed side-by-side for comparable GELYTACs and control constructs. Retroviral supernatant was harvested 48-hr post transfection. Supernatant from replicate dishes were pooled, centrifuged to deplete cell debris, and stored at -80C until use.

### T cell activation

At Day 0, primary human T cells were thawed and activated with anti-CD3/CD28 Human T-Expander Dynabeads (Thermo Fisher) at a 3:1 bead to cell ratio. On Day 2 virus coated culture plates were prepared on non TC-treated 12-well plates that had been pre-coated with RetroNectin (Takara Bio) according to the manufacturer’s instructions, by incubating with 1mL of retroviral supernatant (2x10^7^-5x10^7^ TU/mL) and centrifugation at 3200 RPM, 32 °C for two hrs. The supernatant was subsequently aspirated from the wells and 0.5x10^6^ T cells were added in 1mL of T cell media comprised of: AIM V (Thermo Fisher), 5% fetal bovine serum (FBS), 100 U/mL penicillin (Gibco), 100 mg/mL streptomycin (Gibco), 2 mM L-glutamine (Gibco), 10 mM HEPES (Gibco), and 100 U/mL rhIL2 (Peprotech). After addition of the T cells, the plates were gently spun down at 1200 RPM for 2 min then incubated for 24hrs at 37 °C 5% CO_2_. This transduction process was repeated at Day 3. Dynabeads were removed on Day 4 by magnetic separation. Cells were maintained between 0.4 - 2x10^6^ cells/mL and expanded until Day 10.

### T cell coculture with K562 cells

Coculture experiments were conducted with primary T cells 7-14 days post retroviral transduction. For mCherry and Il6R GELYTACs, 0.6e6 T cells were cocultured with 0.3e6 K562’s. For TGF-β GELYTACs, 0.9e6 T cells were cocultured with 0.1e6 K562 cells. Coculture was set up with complete RPMI supplemented with 100 U/mL rhIL2.

At the time of analysis, the GFP K562’s and T cells were analyzed by flow cytometry for median fluorescence of mCherry or AlexaFluor-647 tagged proteins. GFP and/ or differences in forward and sider scatter were used to distinguish between K562 cells and T cells.

Flow cytometry:

300uL of K562 and or T cell culture was transferred to 96-well V-bottom plate and spun down (500g for 1min). Cells were then washed 1x PBS w/ 0.5% supplemented bovine serum albumen (BSA). Cells were then incubated with SYTOX Blue (1:1000 dilution) for 5 min. Flow cytometry was performed a BioRad ZE5 flow cytometer, and analysis was performed using the FlowJo software package. Gating was performed on single cells and live cells. This instrument is equipped with a 405-nm violet laser, a 488-nm blue laser, a 561-nm green laser and a 639-nm red laser.

For recombinant and HEK293T/ K562 coculture experiments where mCherry was the target protein, K562 cells were washed with PBS then incubated with 5% trypsin for 1 min at 37 °C. After trypsin treatment, trypsin was quenched with PBS supplemented with 0.5% BSA and then subsequently subjected to the standard flow protocol described above. This was done to cleave GELYTACs and receptors on the cell surface that may contribute to mCherry signal from non-internalized mCherry.

### Western blot protocols

Conditioned media from transfected HEK293T cells or transduced T cells was collected (cells were removed by centrifugation) and LDS Sample Buffer (4X) (for reduced blot 4x loading buffer was supplemented with 10% 1M DTT) and boiled at 95 °C for 10 min. 30uL of each sample was separated by SDS–PAGE (10% Bis-Tris gel), then transferred to a nitrocellulose membrane. After transfer, the blot was blocked with Odyssey Blocking Buffer (PBS or TBS) (LI-COR) for 1 h at room temperature with gentle shaking. Membranes were stained with M2 anti-FLAG (Sigma Aldrich) for 1 hr at room temperature or at 4 °C with gentle shaking, then washed three times with PBS-T for five minutes each. The membrane was then incubated with 800CW goat anti-mouse IgG (1:10,000) in Odyssey Blocking Buffer (PBS or TBS) for 1 h at room temperature with gentle shaking. Membranes were washed three times with TBS-T, then imaged using an OdysseyCLxImager (LI-COR). Quantification of band intensities was performed using Image Studio Software (LI-COR).

### Confocal Imagining

K562 cells were plated on HFN coated glass cover slip bottom well plates with phenol red free complete RPMI. On the next day cells were treated with 275 nM of recombinant mCherry GELYTAC or controls and 100 nM of mCherry for 1 hr. The well plates were then imaged live using a Zeiss AxioObserver microscope. This microscope is equipped with 603 oil immersion objectives, outfitted with a Yokogawa spinning disk confocal head, Cascade II:512 camera, a Quad-band notch dichroic mirror (405/488/ 568/647), and 405 (diode), 491 (DPSS), 561 (DPSS) and 640 nm (diode) lasers (all 50 mW). DAPI (405 laser excitation, 445/40 emission), Alexa Fluor488 (491 laser excitation, 528/38 emission) and AlexaFluor647 (640 laser excitation, 700/75 emission) and differential interference contrast (DIC) images were acquired through a 60x oil-immersion lens. Acquisition times ranged from 100 to 2,000 ms. All images were collected and processed using SlideBook 6.0 software (Intelligent Imaging Innovations).

### Knock-out cell line generation

Knock-out K562 cell lines were generated by electroporating the sgRNA (Synthego) Cas9 (ID Technologies) complex using a Lonza 4D Nucleofector. The cell line was generated using Synthego’s protocols.

### Top-Down Protein Mass Spectrometry

After protein expression and purification, the protein samples were further buffer exchanged using a methanol/chloroform/water precipitation and resolubilization method (49). Here, 300 μL of 10 m*M* TCEP prepared in cold LC/MS-grade water (4 °C) was added to 100 μL of protein solution. Then, 400 μL of cold methanol (−20 °C) was added to the protein solution and vortexed for 30 s followed by 100 μL of cold chloroform (−20 °C) and an additional 30 s of vortexing. The sample was centrifuged for 10 min at 18,000*g* at 4 °C after which a biphasic mixture was created with a protein pellet present at the interface. The top layer of the solution was discarded without disturbing the protein pellet. Then, 400 μL of cold methanol (−20 °C) was added to the sample and gently vortexed. The sample was centrifuged for 10 min at 18,000 *g* at 4 °C after which the supernatant was discarded. The cold methanol and centrifugation washing step was repeated two additional times. Protein pellets were resolubilized with 4 μL of 80% formic acid (−20 °C) and diluted to 1% formic acid with 80:20 water: acetonitrile. Top-down LC-MS was carried out using an Agilent 1260 Infinity II high performance liquid chromatography (HPLC) system coupled to an Agilent 6230 ToF LC/MS (Agilent Technologies). Samples were injected onto an Agilent PLRP-S column (2.1 × 50 mm, 5 μm particle size, 1,000 Å pore size) using a gradient of 10% to 90% mobile phase B (0–5 min, 10% B; 5–15 min, 20%–65% B; 15–18 min, 65%–90% B, 18–22 min, 90% B; 22–25 min, 10% B; mobile phase A set to 0.2% formic acid in water; mobile phase B set to 0.2% formic acid in acetonitrile). Flow rate was set to 200 μL/min with a column temperature of 60 °C. Mass spectra were taken at a scan rate of 1 Hz over a 200–3200 *m/z* scan range with the ToF set to extended dynamic range. The mass spectrometer was operated using a dual Agilent jet stream (AJS) high-sensitivity ion source with the following instrument parameters: gas temperature (275°C), drying gas (12 L/min), nebulizer (40 psi), sheath gas temperature (400 °C), sheath gas flow (12 L/min), VCap(3000 V), nozzle voltage (2000 V), fragmentor (250 V), skimmer (65 V), and Oct 1 RF Vpp (750 V). Mass spectra were output from the MassHunter (Agilent Technologies) software and analyzed using MASH Native (50) and UniDec (51).

## Acknowledgments

We thank Kang Yong Loh, Mariko Morimoto, Nicholas Kalogriopoulos, Nicholas Till, and Brendan Floyd for helpful discussions. Figures 4A, 5B, and 5C were created using BioRender.com.

## Funding

This work was supported, in part, by a National Institutes of Health (NIH) grant R01GM058867, OT2CA278686 (C.R.B.), CRUK CGATF-2021/100005 (C.R.B.), by the Chan Zuckerberg Biohub – San Francisco (A. Y. T.) and the Emerson Collective (A. Y. T.). JLY is supported by a BioX SIGF fellowship. C.L.M. and L.L are members of the Parker Institute for Cancer Immunotherapy, which supports the Stanford University Cancer Immunotherapy Program. C.L.M. acknowledges support from the Virginia and D.K. Ludwig Fund for Cancer Research.

## References

1. M. Békés, D. R. Langley, C. M. Crews, PROTAC targeted protein degraders: past is prologue. Nat. Rev. Drug Discov. 21(3), 181–200 (2022).

2. S.M. Banik, K. Pedram, S. Wisnovsky, G. Ahn, N. M. Riley, C. R. Bertozzi, (2020). Lysosome-targeting chimaeras for degradation of extracellular proteins. Nature, 584(7820), 291–297.

3. G. Ahn, S.M. Banik, Miller, C. L., Riley, N. M., Cochran, J. R., C. R. Bertozzi, LYTACs that engage the asialoglycoprotein receptor for targeted protein degradation. Nat. Chem. Biol. 17(9), 937–946 (2021).

4. G. Ahn, N.R, Riley, R.A. Kamber, S. Wisnovsky, S.M. Von Hass, M.C. Bassik, S.M. Banik, C.R. Bertozzi, Elucidating the cellular determinants of targeted membrane protein degradation by lysosome-targeting chimeras. Science. 281 (2023).

5. A.D. Cotton, D.P. Nguyen, J.A. Gramespacher, I. B. Seiple, J. A. Wells, Development of Antibody-Based PROTACs for the Degradation of the Cell-Surface Immune Checkpoint Protein PD-L1. J. Am. Chem. Soc. 143(2), 593–598 (2021).

6. H. Marei, W. T. K. Tsai, Y. S. Kee, K. Ruiz, J. He, C. Cox, T. Sun, S. Penikalapati, P. Dwivedi, M. Choi, D. Kan, P. Saenz-Lopez, K. Dorighi, P. Zhang, Y. T. Kschonsak, N. Kljavin, D. Amin, I. Kim, A. G. Mancini, T. Nguyen, C. Wang, E, Janezic, A. Doan, E. Mai, H. Xi, C. Gu, M. Heinlein, B. Biehs, J. Wu, I. Lehoux, S. Harris, L. Comps-Agrar, D. Seshasayee, F. J. de Sauvage, M. Grimmer, J. Li, N. J. Agard, Chunling Wang, Eric Janezic, Alexander Doan, Elaine Mai, Hongkang Xi, Chen Gu, Melanie Heinlein, Brian Biehs, Jia Wu, Isabelle Lehoux, Seth Harris, Laetitia Comps-Agrar, Dhaya Seshasayee, Frederic J. de Sauvage, Matthew Grimmer, Jing Li, Nicholas J. Agard, F. de Sousa e Melo, F. Antibody targeting of E3 ubiquitin ligases for receptor degradation. Nature, 610 (7930), 182–189 (2022).

7. K. Pance, J. A. Gramespacher, J. R. Byrnes, F. Salangsang, J. A. C. Serrano, A. D. Cotton, V. Steri, J. A. Wells, Modular cytokine receptor-targeting chimeras for targeted degradation of cell surface and extracellular proteins. Nature Biotechnology, 41(2), 273–281, (2023).

8. D. F. Caianiello, M. Zhang, J. D. Ray, R. A. Howell, J. C. Swartzel, E. M. J. Branham, E. Chirkin, V. R. Sabbasani, A. Z. Gong, D. M. McDonald, V. Muthusamy, D. A. Spiegel. Bifunctional small molecules that mediate the degradation of extracellular proteins. Nat. Chem. Biol, 17(9), 947–953 (2021).

9. H. Zhang, Y. Han, Y. Yang, F. Lin, K. Li, L. Kong, H. Liu, Y. Dang, J. Lin, P. R. Chen, Covalently Engineered Nanobody Chimeras for Targeted Membrane Protein Degradation. J. Am. Chem. Soc., 143(40), 16377–16382 (2021).

10. Y. Miao, Q. Gao, M. Mao, C. Zhang, L. Yang, L. Y. Yang D. Han Bispecific Aptamer Chimeras Enable Targeted Protein Degradation on Cell Membranes. Angew. Chem., Int. Ed., 60(20), 11267–11271 (2021).

11. J. Zheng, W. He, J. Li, X. Feng, Y. Li, B. Cheng, Y. Zhou, M. Li, K. Liu, X. Shao, J. Zhang, H. Li, L. Chen, L. Fang, Bifunctional Compounds as Molecular Degraders for Integrin-Facilitated Targeted Protein Degradation. J. Am. Chem. Soc., 144(48), 21831–21836 (2022).

12. R. Sun, Z. Meng, H. Lee, R. Offringa, C. Niehrs. ROTACs leverage signaling-incompetent R-spondin for targeted protein degradation. Cell Chem. Biol., 30(7), 739–752 (2023).

13. E. Loppinet, H. A. Besser, C. E. Lee, W. Zhang, B. Cui, C. Khosla, Targeted Lysosomal Degradation of Secreted and Cell Surface Proteins through the LRP-1 Pathway. J. Am. Chem. Soc., 1–6 (2023).

14. J. Duque-Jimenez, G. Brixi, F. Facchinetti,K. Rhee, W. W. Feng, P. A. Jänne. X. Zhou, Transferrin Receptor Targeting Chimeras (TransTACs) for Membrane Protein Degradation, BioRxiv doi: 10.1101/2023.08.10.552782 (2023).

15. D. H. Siepe, L. K. Picton, & Garcia, K. C. Receptor Elimination by E3 Ubiquitin Ligase Recruitment (REULR): A Targeted Protein Degradation Toolbox. ACS Synth. Biol., 12(4), 1081–1093 (2023).

16. Y. Zhou, P. Teng, N. T. Montgomery, X. Li, W. Tang, Development of Triantennary N-Acetylgalactosamine Conjugates as Degraders for Extracellular Proteins. ACS Cent. Sci., 7(3), 499–506 (2021).

17. X. Zhang, H. Liu, J. He, C. Ou, T. C. Donahue, M. M. Muthana, L. Su, L. X. Wang, Site-Specific Chemoenzymatic Conjugation of High-Affinity M6P Glycan Ligands to Antibodies for Targeted Protein Degradation. ACS Chem. Biol., 17(11),3013–3023 (2022).

18. Y. Wu, B. Lin, Y. Lu, L. Li, K. Deng, S. Zhang, H. Zhang, C. Yang, Z. Zhu, Aptamer-LYTACs for Targeted Degradation of Extracellular and Membrane Proteins. Angew. Chem., Int. Ed., 62(15) (2023).

19. K. Hamada, T. Hashimoto, R. Iwashita, Y. Yamada, Y. Kikkawa, M. Nomizu, Development of a bispecific DNA-aptamer-based lysosome-targeting chimera for HER2 protein degradation. Cell Rep. Phys. Sci., 4(3), 101296 (2023)

20. R. A. Howell, R. Wang, D. M. Mcdonald, D. A. Spiegel, Bifunctional molecules that induce targeted degradation and transcytosis of extracellular proteins in brain cells, ChemRxiv doi:10.26434/chemrxiv-2023-5l14b-v2, (2023).

21. B. Huang, M. Abedi, G. Ahn, B. Coventry, I. Sappington, A. Chazin-gray, S. Chan, S. Gerben, A. Murray, S. Wang, J. O’Neill, N. R. Bennett, P. Venkatesh, D. Satoe, M. Ahlrichs, C. Dobbins, W. Yang, X. Wang, D. Vafeados, R. Mout, S. Shivaei, L. Cao, L. Carter, L. Stewart, J. B. Spangler, G. J. L. Bernardes, K. T. Roybal, P. Jr. Greisen, X. Li, C. Bertozzi, David Baker, Designed Endocytosis-Triggering Proteins mediate Targeted Degradation. BioRxiv [Preprint] (2023). Doi: 10.1101/2023.08.19.55332.

22. B. Zhang, R. K. Brahma, L. Zhu, J. Feng, S. Hu, L. Qian S. Du Insulin-like Growth Factor 2 (IGF2)-Fused Lysosomal Targeting Chimeras for Degradation of Extracellular and Membrane Proteins, J. Am. Chem. Soc. XXXX, XXX, XXX−XXX (2023).

23. L. Labanieh, C. L. Mackall, CAR immune cells: design principles, resistance and the next generation. Nature, 614(7949), 635–648 (2023).

24. B. D. Choi, X. Yu, A. P. Castano, A. A. Bouffard, A. Schmidts, R. C. Larson, S. R. Bailey, A. C. Boroughs, M. J. Frigault, M. B. Leick, I. Scarfò, C. L. Cetrulo, S. Demehri, B. V. Nahed, D. P. Cahill, H. Wakimoto, W. T. Curry, B. S. Carter, M. V. Maus, CAR-T cells secreting BiTEs circumvent antigen escape without detectable toxicity. Nat. Biotechnol. 37(9), 1049–1058 (2019).

25. Y. Yin, J. L. Rodriguez, N. Li, R. Thokala, M. P. Nasrallah, L. Hu, L. Zhang, J. V. Zhang, M. T. Logun, D. Kainth, L. Haddad, Y. Zhao, T. Wu, E. X. Johns, Y. Long, H. Liang, J. Qi, X. Zhang, Z. A. Binder, Z. Lin, D. M. O’Rourke. Locally secreted BiTEs complement CAR T cells by enhancing killing of antigen heterogeneous solid tumors. Mol. Ther., 30(7), 2537–2553, (2022).

26. S. Rafiq, O. O. Yeku, H. J. Jackson, T. J. Purdon, D. G. van Leeuwen, D. J. Drakes, M. Song, M. M. Miele, Z. Li, P. Wang, S. Yan, J. Xiang, X. Ma, V. E. Seshan, R. C. Hendrickson, C. Liu, R. J. Brentjens. Targeted delivery of a PD-1-blocking scFV by CAR-T cells enhances anti-tumor efficacy in vivo. Nat. Biotechnol., 36(9), 847–858 (2018).

27. W. Xia, J. Chen, W. Hou, J. Chen, Y. Xiong, H. Li, X. Qi, H. Xu, Z. Xie, M. Li, X. Zhang, J. Li. Engineering a HER2-CAR-NK Cell Secreting Soluble Programmed Cell Death Protein with Superior Antitumor Efficacy. Int. J. Mol. Sci., 24(7), 1–17 (2023).

28. K. J. Curran, B. A. Seinstra, Y. Nikhamin, R. Yeh, Y. Usachenko, D. G. Van Leeuwen, T. Purdon, H. J. Pegram, R. J. Brentjens, Enhancing antitumor efficacy of chimeric antigen receptor T cells through constitutive CD40L expression. Mol. Ther. 23(4), 769–778 (2015).

29. O. O. Yeku, T. J. Purdon, M. Koneru, D. Spriggs, R. J. Brentjens, Armored CAR T cells enhance antitumor efficacy and overcome the tumor microenvironment. Sci. Rep., 7(1), 1–14 (2017).

30. H. J. Pegram, J. C. Lee, E. G. Hayman, G. H. Imperato, T. F. Tedder, M. Sadelain, R. J. Brentjens, Tumor-targeted T cells modified to secrete IL-12 eradicate systemic tumors without need for prior conditioning. Blood, 119(18), 4133–4141 (2012).

31. J. Brown, C. Delaine, O. J. Zacheo, C. Siebold, R. J. Gilbert, G. Van Boxel, A. Denley, J. C. Wallace, A. B. Hassan, B. E. Forbes, E. Y. Jones. Structure and functional analysis of the IGF-II/IGF2R interaction. EMBO, 27(1), 265–276 (2008).

32. Y. Oka, L. M. Rozek, M. P. Czech, Direct demonstration of rapid insulin-like growth factor II receptor internalization and recycling in rat adipocytes. Insulin stimulates 125I-insulin-like growth factor II degradation by modulating the IGF-II receptor recycling process. JBC, 260(16), 9435–9442 (1985).

33. M. E. Zavorka, C.M. Connelly, R. Grosely, G. Richard McDonald Oncotarget Inhibition of insulin-like growth factor II (IGF-II) -dependent cell growth by multidentate pentamannosyl 6-phosphate-based ligands targeting the mannose 6-phosphate / IGF-II receptor. 7(38), 10–16, (2016).

34. P. C. Fridy, Y. Li, S. Keegan, M. K. Thompson, I. Nudelman, J. F. Scheid, M. Oeffinger, M. C. Nussenzweig, D. Fenyö, B. T. Chait, M. P. Rout, A robust pipeline for rapid production of versatilenanobody repertoires. Nat. Methods, 11(12), 1253–1260 (2014).

35. L. Huang, D. Pike, D.E. Sleat, V. Nanda, P. Lobel, Potential pitfalls and solutions for use of fluorescent fusion proteins to study the lysosome. PLoS ONE, 9(2). (2014).

36. C. Delaine, C. L. Alvino, K. A. McNeil, T. D. Mulhern, L. Gauguin, P. De Meyts, E. Y. Jones, J. Brown, J. C. Wallace, B. E. Forbes. A novel binding site for the human insulin-like growth factor-II (IGF-II)/mannose 6-phosphate receptor on IGF-II. JBC, 282(26), 18886–18894 (2007).

37. S. S. Lam, J. D. Martell, K. J. Kamer, T. J. Deerinck, M. H. Ellisman, V. K. Mootha, A. Y. Ting Directed evolution of APEX2 for electron microscopy and proximity labeling. Nat. Methods, 12(1), 51–54 (2014).

38. T. C. Branon, J. A. Bosch, A. D. Sanchez, N. D. Udeshi, T. Svinkina, S. A. Carr, J. L. Feldman, Perrimon, N., Ting, A. Y, Efficient proximity labeling in living cells and organisms with TurboID. Nat. Biotechnol., 36(9), 880–898 (2018).

39. J. D. Martell, M. Yamagata, T. J. Deerinck, S. Phan, C. G. Kwa, M. H. Ellisman, J. R. Sanes, A. Y. Ting, A split horseradish peroxidase for the detection of intercellular protein-protein interactions and sensitive visualization of synapses. Nat. Biotechnol., 34(7), 774–780 (2016).

40. S. Y. Lee, J. S. Cheah, B. Zhao, C. Xu, H. Roh, C. K. Kim, K. F. Cho, N. D. Udeshi, S. A. Carr, A. Y. Ting, Engineered allostery in light-regulated LOV-Turbo enables precise spatiotemporal control of proximity labeling in living cells. Nat. Methods, 20(6), 908–917 (2023).

41. C. K Ala, A. L. Klein, J.J. Moslehi. Cancer Treatment-Associated Pericardial Disease: Epidemiology, Clinical Presentation, Diagnosis, and Management. Curr. Cardiol. Rep. 21(12), 1–9 (2019).

42. A. Moulin, M. Mathieu, C. Lawrence, R. Bigelow, M. Levine, C. Hamel, J. P Marquette, J. Le Parc, C. Loux, P. Ferrari, C. Capdevila, J. Dumas, B. Dumas, A. Rak, J. Bird, H. Qiu, C. Q. Pan, T. Edmunds, T. R. R. Wei. Structures of a pan-specific antagonist antibody complexed to different isoforms of TGFβ reveal structural plasticity of antibody-antigen interactions. Protein Sci., 23(12), 1698–1707 (2014).

43. M. Van Roy, C. Ververken, E. Beirnaert, S. Hoefman, J. Kolkman, M. Vierboom, E. Breedveld, B. ‘t Hart, S. Poelmans, L Bontinck, A. Hemeryck, S. Jacobs, S. J. Baumeister, H. Ulrichts. (2015). The preclinical pharmacology of the high affinity anti-IL-6R Nanobody® ALX-0061 supports its clinical development in rheumatoid arthritis. Arthritis Res. Ther., 17(1) (2015).

44. A. S. De Groot, D. W. Scott Immunogenicity of protein therapeutics. Trends Immunol 28(11):482–490 (2007).

45. H. Schellekens Bioequivalence and the immunogenicity of biopharmaceuticals. Nat Rev Drug Discov, 1(6):457–462 (2002).

46. Lu, H., Cheng, Z., Hu, Y., & Tang, L. V. What Can De Novo Protein Design Bring to the Treatment of Hematological Disorders? Biology, 12(2). (2023).

47. Shah, D. K. Pharmacokinetic and pharmacodynamic considerations for the next generation protein therapeutics. J. Pharmacokinet. Pharmacodyn, 42(5), 553–571 (2015).

48. H. T. Rogers, D. S. Roberts, E. J. Larson, J. A. Melby, K. J. Rossler, A. V. Carr, K. A. Brown, and Y. Ge, Comprehensive Characterization of Endogenous Phospholamban Proteoforms Enabled by Photocleavable Surfactant and Top-down Proteomics, Anal. Chem., 95 (35), 13091–13100 (2023).

49. E.J. Larson, M. R. Pergande, M. E. Moss, K. J. Rossler, R. K. Wenger, B. Krichel, H. Josyer, J. A. Melby, D. S. Roberts, K. Pike, Z. Shi, H. J. Chan, B. Knight, H. T. Rogers, K. A. Brown, I. M. Ong, K. Jeong, M. T. Marty, S. J. McIlwain, Y. Ge, MASH Native: a unified solution for native top-down proteomics data processing, Bioinformatics, 39 (6), (2023).

50. M. T. Marty, A. J. Baldwin, E. G. Marklund, G. K. A. Hochberg, J. L. P. Benesch, C. V. Robinson, Bayesian Deconvolution of Mass and Ion Mobility Spectra: From Binary Interactions to Polydisperse Ensembles, Anal. Chem., 87 (8), 4370–4376 (2015).

