## Supporting Information for "Directed Evolution of Genetically Encoded LYTACs for Cell-Mediated Delivery"

**This PDF file includes:**

Supporting text
Figures S1 to S6
Tables S1 to S5
SI References

**Supporting Information Text**

Yeast cell culture.

*S. cerevisiae* strain EBY100 was cultured according to previously published protocols (1, 2, 3, 4). Cells were propagated at 30 °C in synthetic dextrose plus casein amino acid (SDCAA, 'regular') medium supplemented with tryptophan (20 mg/L). Yeast cells were transformed with the yeast-display plasmid pCTCON2 using the Frozen E-Z Yeast Transformation II kit (Zymoprep) according to manufacturer protocols. Transformed cells containing the *TRP1* gene were selected on SDCAA plates and propagated in SDCAA medium at 30 °C. Protein expression was induced by inoculating saturated yeast culture into SGCAA (synthetic galactose plus casein amino acid), at a 1:10 dilution and incubating at 30 °C for 18 – 24 h.

Generation of ligase libraries for yeast display.

Libraries of ligase mutants were generated by error-prone PCR according to published protocols (ref 41). 150 ng of the template ligase in vector pCTCON2 was amplified for 15 rounds with 0.4 μM forward and reverse primers:

F: 5′- GGA CAA GGA ACC CAG GTG ACC GTG TCA TCT CCG TTT ACC - 3′

R: 5′- CGA TTT TGT TAC ATC TAC ACT GTT GTT ATC AGA TCT CGA G -3′

and 2 mM $MgCl_2$, 5 units of *Taq* polymerase (NEB), and 20 μM of mutagenic nucleotide analogs 8-oxo-2′-deoxyguanosine-5′-triphosphate (8-oxo-dGTP) and 0.5uM-20uM 2′-deoxy-P-nucleoside-5′-triphosphate (dPTP). The PCR products were then gel purified and reamplified for another 30 cycles under normal PCR conditions using the same primers.

The inserts were then electroporated into electrocompetent *S. cerevisiae* EBY100 with the BamHI-XhoI linearized pCTCON2 vector (4 μg mutinagenized insert/1 μg cut vector) backbone. The electroporated cultures were rescued in 2 mL yeast extract peptone dextrose (YPD) complete medium for 1 h at 30 °C with no shaking. Cells were vortexed briefly, and 1.99 mL of the rescued cell suspension was transferred to 100 mL of SDCAA medium supplemented with 50 units/mL penicillin, 50 μg/mL streptomycin, and 50ug/mL kanamycin and grown for 2 days at 30 °C. The remaining 10 μL of the rescued cell suspension was diluted 100×, 1000×, 10000×, and 100000×; 20 μL of each dilution was plated on SDCAA plates and incubated at 30 °C for 3 days. After 3 days, each colony observed in the 100×, 1000×, 10000×, or 100000× segments of plates will correspond to $10^4$, $10^5$, $10^6$, or $10^7$ transformants in the library, respectively.

General methods for yeast display-based directed evolution.

For each round of evolution, we input 10-fold more yeast cells than the estimated library size. For the first round, library size was estimated by the transformation efficiency of the initial ligase library. For subsequent rounds, library size was taken to be the number of yeast cells collected during the previous sort. GELYTACs protein expression was induced by inoculating saturated yeast culture in SDCAA into SGCAA (1:10 dilution) and incubating at 30 °C for 18 – 24 h.

For GELYTACs and IGF2R binding, yeast cells were then incubated with 100 uL (for analysis) or 3mL (for sorting) PBS-B (2% Bovine Serum Albumin in PBS) with 100nM of either biotinylated Fc fused IGF2R D11-13 (GP Biosciences, used for library #1) or Alexa-Fluor 647 or 488 labeled IGF2R D11-13 (R&D systems). These reagents were stained for 1hr over 4º C; for biotinylated Fc fused IGF2R, the samples were washed 3x with PBS-B and then stained with 1:50

streptavidin- phycoerythrin (PE) in PBS-B. GELYTACs expression was monitored with an anti myc antibody (either mouse fluorescently labeled anti-myc clone 9E10 from R&D or chicken anti-myc). For chicken ani-myc antibody, an Alexa Fluor-488 labeled anti-chicken goat. For the final sort for library #1, we performed a competitive wash after IGF2R labeling by incubating labeled yeast cultures with 1uM IGF2 in PBS-B for 1 hr at 37$^{o}$C.

For two-dimensional FACS sorting, samples were resuspended in PBS-B at a maximal concentration of 100 million cells/mL and sorted on a BD FACS Aria II cell sorter (BD Biosciences) with the appropriate lasers and emission filters (488 nm excitation laser and 525/50 emission filter, 561 nm excitation laser and 582/15 emission filter for PE, 640 nm excitation laser and 670/30 emission filter for AF647). To analyze and sort single yeast cells, cells were plotted by a forward-scatter area (FSC-A) and side-scatter area (SSC-A) and a gate was drawn around cells clustered between $10^2$ – $10^4$ FSC-A, 30k – 150K SSC-A to give population P1. Cells from population P1 were then plotted by FSC-A and forward-scatter height (FSC-H) and a gate was drawn around cells clustered between 20K-110K FSC-A, 10K-90K FSC-H to give population P2. Cells from population P2 were then plotted by forward-scatter width (FSC-W) and forward-scatter height (FSC-H) and a gate was drawn around cells clustered between 250 – 3K SSC-H and 64K– 75K SSC-W to give population P3. Population P3 often represented ~50%% of the total population analyzed.

From population P3, gates were drawn to collect yeast with the highest activity/expression ratio, i.e., positive for AF647 signal that also had high PE (library #1) or AF488 signal (library #2). After sorting, yeast were collected in SDCAA medium containing 1% penicillin-streptomycin, 1% kanamycin, and 1% chlorophenol and incubated at 30 °C for 24 h. 1 mL of the growing culture was removed for DNA extraction using the Zymoprep yeast Plasmid Miniprep II (Zymo Research) kit according to manufacturer protocols (using 6 μL zymolyase, vigorously vortex after lysis), and at least ten-fold excess of the number of cells retained during sorting were propagated in SDCAA + 1% pen-strep to ensure oversampling (yeast cells were passaged in this manner at least two times prior to the next round of selection). To analyze yeast populations and clones by FACS, yeast samples were prepared on a small scale (1 mL cultures) as described above and analyzed on a BioRad ZE5 flow cytometer (BioiRad). FlowJo software was used to analyze all data from FACS sorting and analysis. Summaries of all yeast-display directed evolution and resulting mutants are shown in (Figure S2A-C), and are described in detail in the 'Directed evolution of GELYTACs' sections below.

Directed evolution of GELYTACs: generation 1.

For the first round of evolution three libraries were generated using BirA-R118S as the starting template. The three libraries were generated using error prone PCR as described above, using the following conditions to produce varying levels of mutagenesis:

Library 1: 2 μM 8-oxo-dGTP, 0.2 μM dPTP, 15 PCR cycles

Library 2: 2 μM 8-oxo-dGTP, 1 μM dPTP, 15 PCR cycles

Library 3: 2 μM 8-oxo-dGTP, 2 μM dPTP, 15 PCR cycles

Each library showed a spread of binding (Figure S2A) The library sizes (approximated by number of transformants as described above). Pooled together, the library size was estimated to be 1.6 x$10^7$. Sequencing of 30 clones of pooled Library 1 revealed an average of 2.1 amino acid changes per GELYTAC gene (supplementary table 1).

Library was passaged twice, induced in SGCAA, and labeled using protocols described above. The top 1.4% of cells ($1.8x10^6$ cells) were sorted by FACS by the gating scheme described above.

Generation #1 Sort #1 was passaged twice and analyzed by FACS side-by-side with the WT to ensure the sort was successful (resulting population still had expression and had higher binding).

Generation #1 Sort #1 was then induced in SGCAA and labeled with IGF2R and washed competitively with IGF2 (described above). The top 0.4% of cells ($2.4x10^5$ cells) according to the gating scheme described previously was sorted.

Sequencing of 30 clones reveled convergence on mutation F19L. Comparison of F19L to the wild type scaffold revealed an increase of 1.94 increase in Alexa 488 IGF2R signal. This mutation was previously reported by Brown et. al to gain affinity (5). After Generation #1 Sort #2, we made subsequent libraries from F19L, along with other known mutations to increase IGF2 affinity for IG2R (G6A and Y27L).

Directed evolution of GELYTACs: generation 2.

The Generation #2 library is composed of the following sub-libraries:

Library 1:  Template: F19L, 2 μM 8-oxo-dGTP, 0.2 μM dPTP, 15 PCR cycles

Size: 1,150,000

Library 2: Template: F19L, 2 μM 8-oxo-dGTP, 1 μM dPTP, 15 PCR cycles

Size: 2400000

Library 3: Template: F19L,  2 μM 8-oxo-dGTP, 2 μM dPTP, 15 PCR cycles

Size: 1300000

Library 4: Template: G6A,  2 μM 8-oxo-dGTP, 0.2 μM dPTP, 15 PCR cycles

Size: 1850000

Library 5: Template: G6A, 2 μM 8-oxo-dGTP, 1 μM dPTP, 15 PCR cycles

Size: 1300000

Library 6: Template: G6A, 2 μM 8-oxo-dGTP, 2 μM dPTP, 15 PCR cycles

Size: 1600000

Library 7: Template: Y27L, 2 μM 8-oxo-dGTP, 0.2 μM dPTP, 15 PCR cycles

Size: 850000

Library 8: Template: Y27L, 2 μM 8-oxo-dGTP, 1 μM dPTP, 15 PCR cycles

Size: 1650000

Library 9: Template: Y27L, 2 μM 8-oxo-dGTP, 2 μM dPTP, 15 PCR cycles

Size: 1750000

These libraries were all analyzed by flow cytometry, and all had good expression and varying degrees of binding to IGF2R (Figure S2B); thus all sub libraries were combined to form Generation #2 combined library and yielded a total size of 1.4 x $10^7$.

Library was passaged twice, induced in SGCAA, and labeled using protocols described above (using monomeric, Alexa Fluor 488). The top 1.5% of cells (8.2x$10^5$ cells) were sorted by FACS by the gating scheme described above. 29 colonies were sequenced and there was an average of 1.1 mutations per construct (Figure S2B, supplementary table 3).

Generation #2 Sort #1 was then induced in SGCAA and labeled with IGF2R (monomeric, Alexa Fluor 488). The top 0.5% of cells according to the gating scheme described previously was sorted. The sorted yeast were then passaged twice and subsequently sorted again using the same gating scheme; the top 0.6% yeast were sorted (114,577 cells). 45 colonies were sequenced averaged 0.9 mutations per construct (Figure S2B, supplementary table 4)

Generation #2 sort#3, induced in SGCAA, and labeled using protocols described above (using monomeric, Alexa Fluor 488). The top 0.6% of cells (8.2x$10^5$ cells) were sorted by FACS by the gating scheme described above. 29 colonies were sequenced and there was an average of 1.1 mutations per construct (Figure S2B, supplementary table 4). The mutations are outlined in supplementary table 4.

After the first sort of Generation #2, the yeast library became infected with an unknown contaminant. To remove the infection, yeast cells were washed with SDCAA supplemented with hydrocholoric acid to pH 2. Low pH conditions have been used to remove infection from yeast cultures.

**Supplemental Figures:**

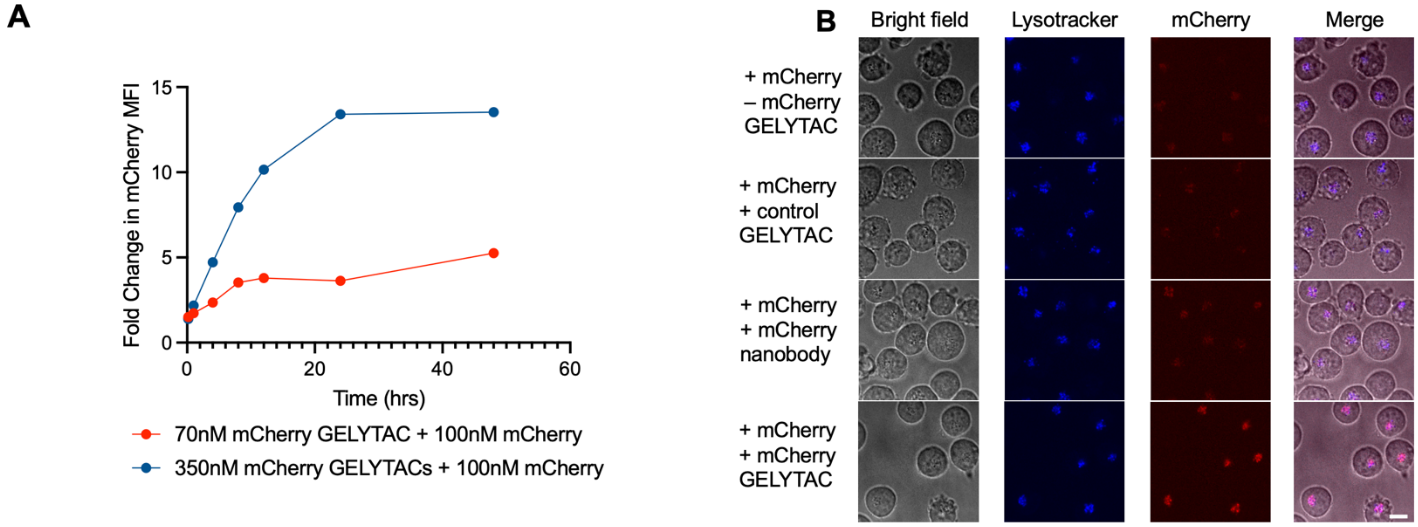


**Supplemental figure 1. Extended Recombinant GELYTAC mediate internalization of mCherry by K562 cells.** A) Time course of fold change of mCherry MFI in K562 cells treated with either 70 nM or 350 nM mCherry GELYTAC. B) Confocal imaging experiment from Figure 2C with the addition of control GELYTAC and mCherry nanobody only controls. Scale bars, 10 μm.

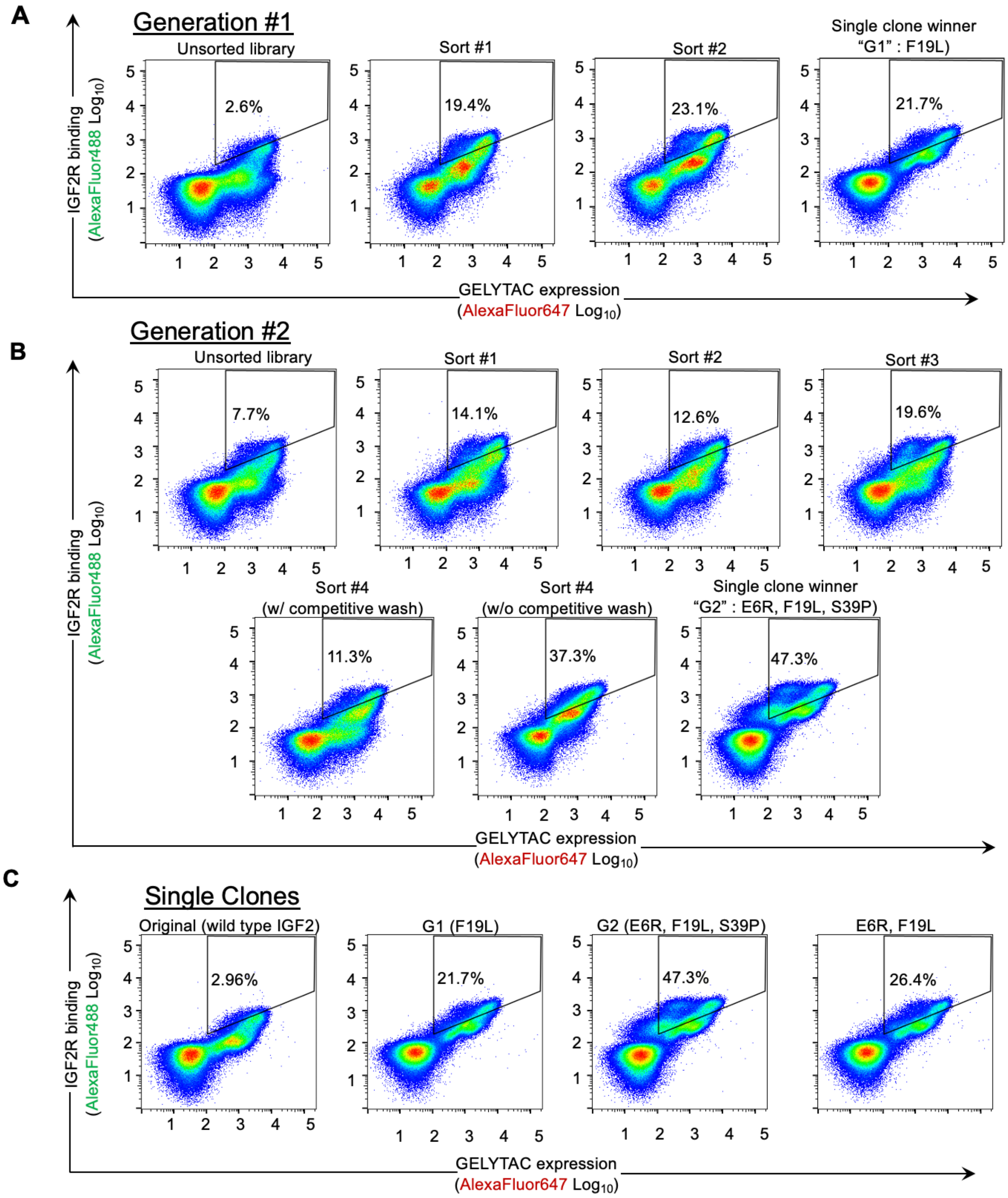


**Supplemental figure 2. Directed evolution progression** A) 2D FACS plots for generation #1's directed evolution from unsorted library to final sort to the winning clone G1 (F19L). B) 2D FACS plots for generation #2's directed evolution from unsorted library to final sort to the winning clone G2 (E6R, F19L, S39P). C) Comparison of single clones (original, G1, G2, and E6R + F19L) showing the increase of GELYTACs binding to IGF2R and that the S39S mutation from evolution is critical in addition to the published E6R and F19L mutations for G2 GELYTAC's improvement.

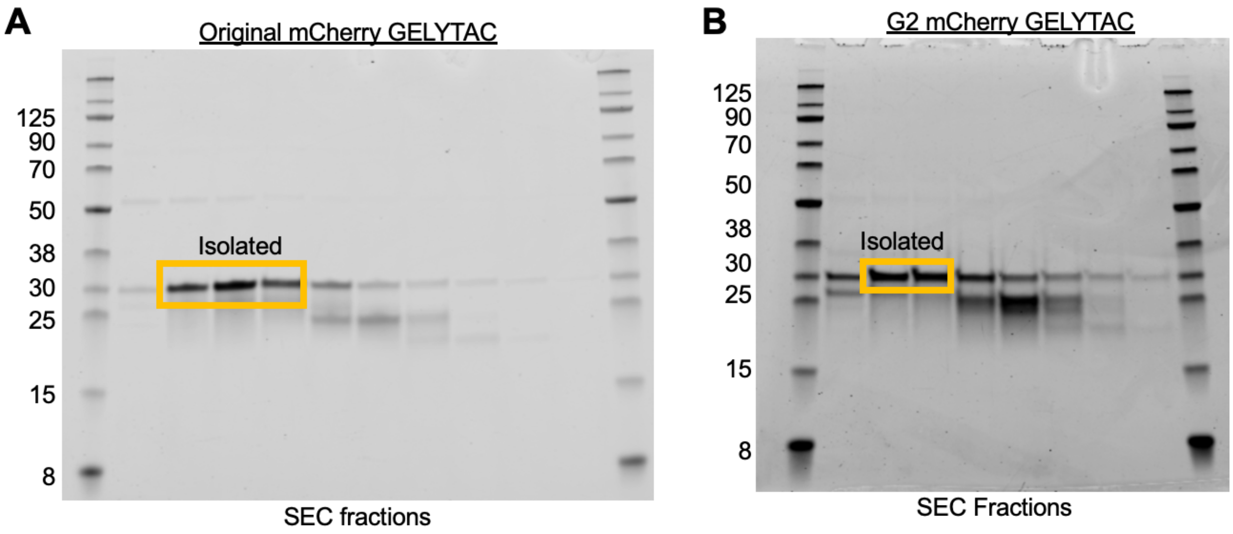


**Supplemental figure 3. Recombinant GELYTAC purification.** A) Coomassie gel of size exclusion chromatography (SEC) fractions from the purification of original mCherry GELYTAC. Only the monomer was isolated and used for experiments. B) Coomassie gel of size exclusion chromatography (SEC) fractions from the purification of G2 mCherry GELYTACs. Only the monomer was isolated and used for experiments.

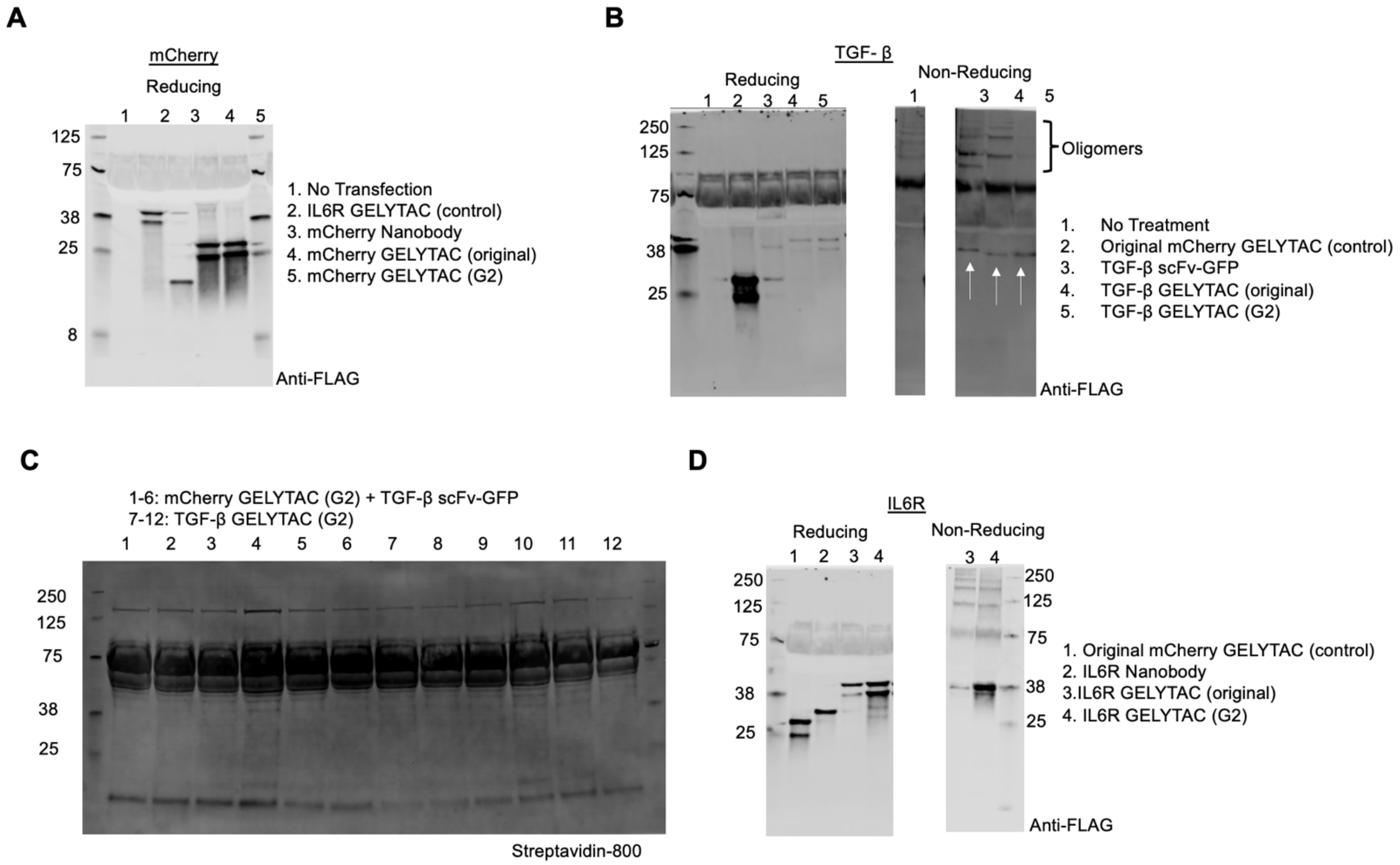


**Supplemental figure 4. Secretion of GELYTAC from HEK293T cells.** A) Anti-FLAG Western blot of supernatant from HEK293T cells secreting mCherry GELYTAC and controls under reducing conditions. B) Anti-FLAG Western blot of supernatant from HEK293T cells secreting TGF-β GELYTACs and controls under reducing and non-reducing conditions. C) Anti-FLAG Western blot of supernatant from HEK293T cells secreting IL6R GELYTACs and controls under reducing and non-reducing conditions. D) Streptavidin-800 western blot of biotinylated TGF- β degradation upon treatment with HEK293T cells secreting both mCherry GELYTAC and TGF-β scFv or TGF-β GELTAC.

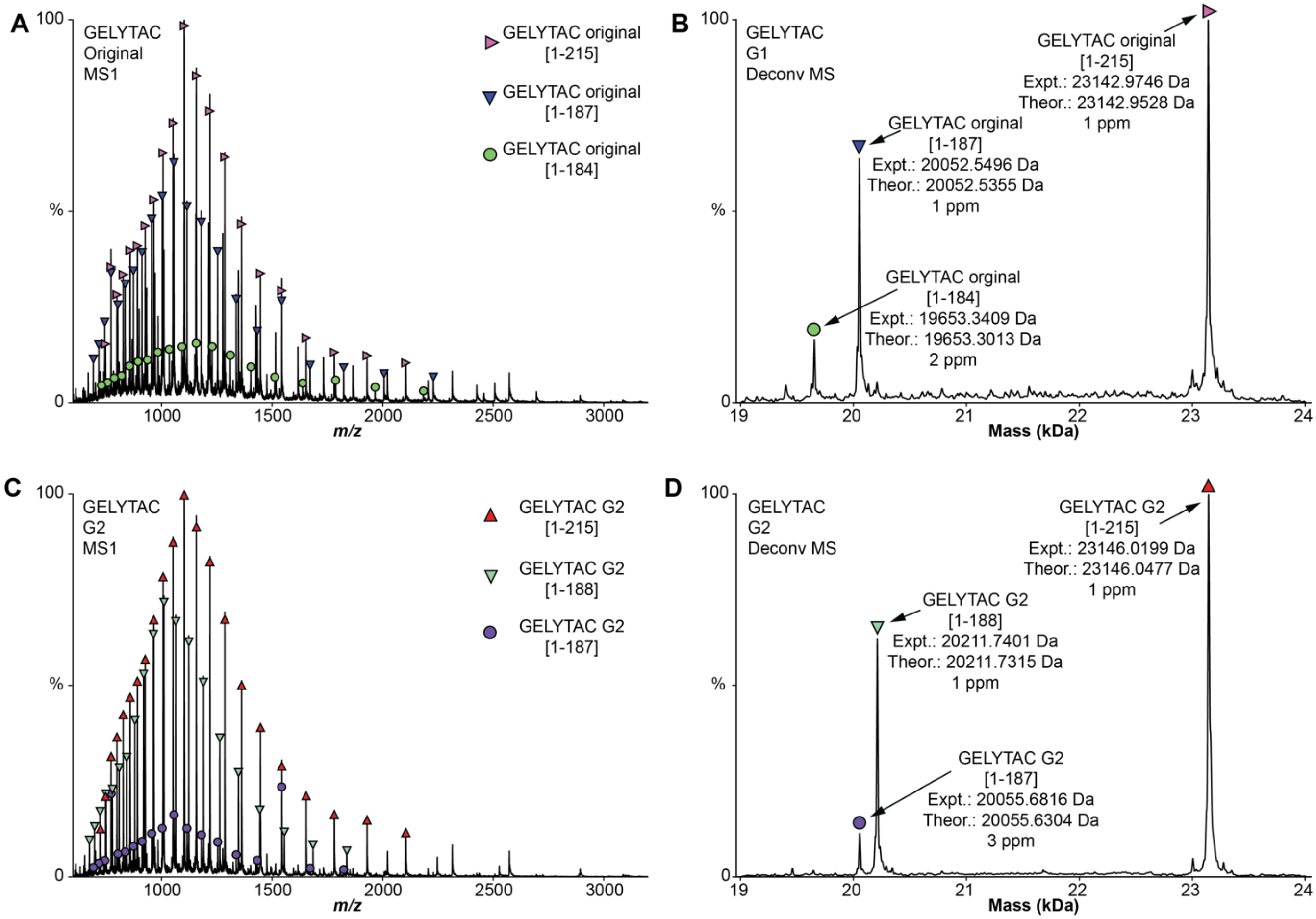


**Supplemental figure 5. Determining cleavage product from mCherry GELYTACs secreted by mammalian cells.** A) Top-down mass spectrometry trace from anti-FLAG pulldown from supernatant of HEK293T cells secreting original mCherry GELYTAC. From the trace, 3 different species are observed. B) Identification of the cleavage products as truncations of original mCherry GELYTAC. C) Top-down mass spectrometry trace from anti-FLAG pulldown from supernatant of HEK293T cells secreting G2 mCherry GELYTAC. From the trace, 3 different species are observed. B) Identification of the cleavage products as truncations of G2 GELYTAC.

A

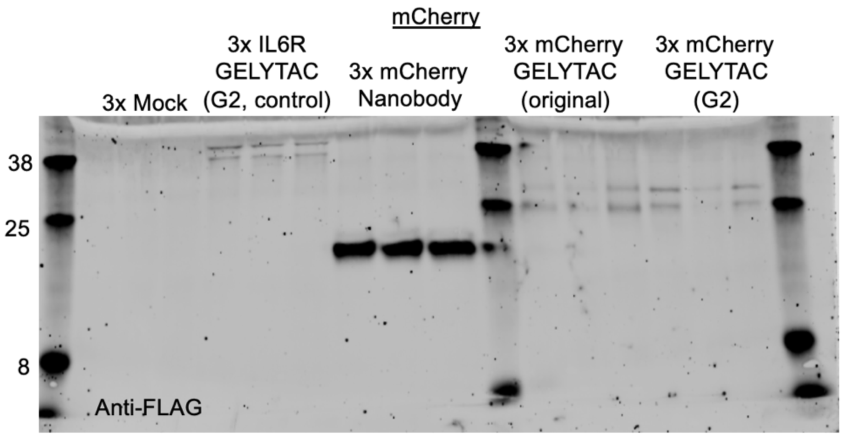


B

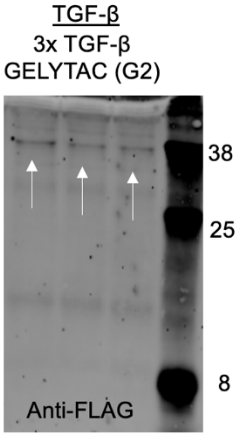


C

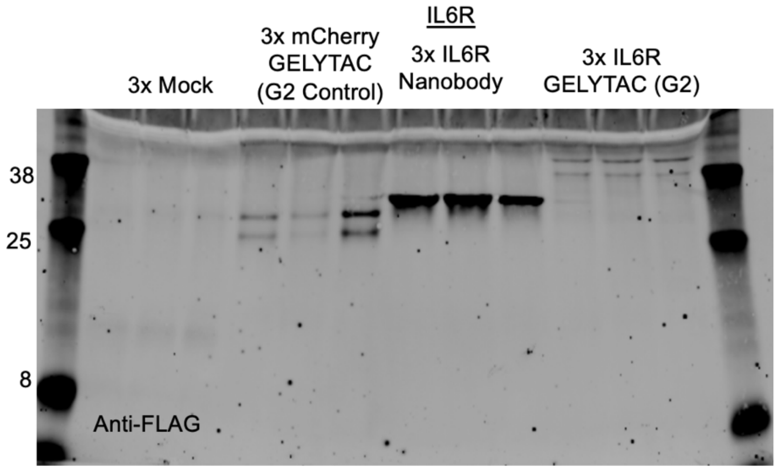


**Supplemental figure 6. Secretion of GELYTAC from T cells.** A) Anti-FLAG Western blot of supernatant from T cells secreting mCherry GELYTACs and controls under reducing conditions. 3 donors shown. B) Anti-FLAG Western blot of supernatant from T cells secreting TGF-β GELYTACs under reducing conditions. 3 donors shown. C) Anti-FLAG Western blot of supernatant from T cells secreting IL6R GELYTACs and controls under reducing conditions. 3 donors shown.

| S5 | E6 | C9 | E12 | D15 | L17 | C21 | D23 | F28 | R34 | G41 | E45 | C47 | F48 | S50 | L53 | T58 | Y59 | T62 |
|---|---|---|---|---|---|---|---|---|---|---|---|---|---|---|---|---|---|---|
| | | | | | | | | | | | | | | | | | | T62P |
| | | | | | | | | | | | | | | | V53A | | | |
| | | | | | | | | | | | | | | S50G | | | | |
| | | | | | | | | | | G41E | | | | | | | | |
| | | | E12G | T16A | | | | F28L | | | | | | | | T58A | Y59H | |
| | | | | | | | | | | | E45G | | | | | | | |
| S5N | E6G | | | | | | | | | | | | | | | | | |
| | | C9Y | E12G | | | | | | | | | | | | | | | |
| | | | | | | | | F28V | | | | | F48L | | | | | |
| | | | | | | | | | | | | C47R | | | | | | |
| | | | | | L17I | | | | | | | | | | | | | |
| | | | | | | | | | V34A | | | | | | | | | |
| | | | | | | | D23G | | | | | | | | | | | |
| | | | | | | C21G | | | | | | | | | | | | |

**Supplemental table 1. Sequencing of unsorted library from Generation #1 (non-template clones).**
**Supplemental table 2. Sequencing after sort #2 from Generation #1 (non-template clones)..**

| R3 | P4 | T7 | F19 |
|---|---|---|---|
| | | | F19L |
| | | | F19L |
| | | | F19L |
| | | | F19L |
| | | | F19L |
| | | | F19L |
| | | | F19L |
| | | | F19L |
| | | | F19L |
| | | | F19L |
| | | | F19L |
| | | | F19L |
| | | | F19L |
| | | | F19L |
| | | | F19L |
| | | | F19L |
| | | | F19L |
| | | | F19L |
| | | | F19L |
| | | | F19L |
| | | | F19L |
| | | | F19L |
| | | | F19L |
| | | | F19L |
| | | | F19L |
| R3S | | | |
| | P4L | T7A | |

Shows convergence on F19L mutation.

| Y2 | E6 | L13 | T16 | F19 | Y27 | F28 | R34 | C47 | C51 | D52 | L53 | T58 | Y59 | A64 |
|---|---|---|---|---|---|---|---|---|---|---|---|---|---|---|
| | E6R | | T16A | | | | | | | | | | | |
| | E6R | | T16A | | | | | | | | | | | |
| | E6R | | T16A | | | | | | | | | | | |
| | E6R | | | | | | | | | | L53P | | | |
| | E6R | | | | | | | | | | L53P | | | |
| | | | | | Y27L | | | C47Y | | | | | | |
| | | | | | Y27L | F28S | | | | | | | | |
| | | | | F19L | | | | | | | | | Y59C | |
| | E6R | | | | | | | | | | | | | A64G |
| | | L13P | | F19I | Y27L | | | | | D52G | | | | |
| | E6R | | | F19L | | | | | | | | | | |
| | | | | | Y27L | | | | C51Y | | | | | |
| Y2C | | | | | Y27L | | | | | | | | | |
| | E6R | | | | | | | | | | | T58G | | |
| | E6R | | | | | | R34G | | | | | | | |

**Supplemental table 3. Sequencing after sort #1 from Generation #2 (non-template clones).**

| E6 | T7 | T16 | F19 | Y27 | S39 | C51 | D52 |
|---|---|---|---|---|---|---|---|
| E6R | | | F19L | | S39P | | |
| E6R | | | F19L | | S39P | | |
| E6R | | | F19L | | S39P | | |
| E6R | | | F19L | | S39P | | |
| E6R | | | F19L | | S39P | | |
| E6R | | | F19L | | S39P | | |
| | | | | Y27L | | C51Y | |
| | | | | Y27L | | C51Y | |
| | | | | Y27L | | C51Y | |
| | | | | Y27L | | C51Y | |
| | | | | Y27L | | C51Y | |
| E6R | T7A | | | | | | |
| E6R | T7A | | | | | | |
| | | | | Y27L | | | D52G |
| E6R | | | | Y27C | | | |
| E6R | | T16A | | | | | |

**Supplemental table 4. Sequencing after sort #3 from Generation #2 (non-template clones)..**

| E6 | F19 | Y27 | S33 | S39 | R40 | A54 | E57 | T62 |
|---|---|---|---|---|---|---|---|---|
| E6R | F19L | | | S39P | | | | |
| E6R | F19L | | | S39P | | | | |
| E6R | F19L | | | S39P | | | | |
| E6R | F19L | | | S39P | | | | |
| E6R | F19L | | | S39P | | | | |
| E6R | F19L | | | S39P | | | | |
| E6R | F19L | | | S39P | | | | |
| E6R | F19L | | | S39P | | | | |
| E6R | F19L | | | S39P | | | | |
| E6R | F19L | | | S39P | | | | |
| E6R | F19L | | | S39P | | | | |
| E6R | F19L | | | S39P | | | | |
| E6R | F19L | | | S39P | | | | |
| E6R | F19L | | | S39P | | | | |
| E6R | F19L | | | S39P | | | | |
| E6R | F19L | | | S39P | | | | |
| E6R | F19L | | | S39P | | | | |
| E6R | F19L | | | S39P | | | | |
| E6R | F19L | | | S39P | | | | |
| E6R | F19L | | | S39P | | | | |
| E6R | F19L | | | S39P | | | | |
| E6R | F19L | | | S39P | | | | |
| E6R | F19L | | | S39P | | | | |
| E6R | F19L | | | S39P | | | | |
| E6R | F19L | | | S39P | | | | |
| E6R | F19L | | | S39P | | | | |
| E6R | F19L | | | S39P | | | | |
| E6R | F19L | | | S39P | | | | |
| E6R | | | | S39P | | | | |
| E6R | | | | | | E57A | | |
| E6R | | | | | R40G | | | |
| E6R | | | | | | | E57G | |
| E6R | | | | | | | | T62A |
| | | Y27L | S33P | | | | | |

**Supplemental table 5. Sequencing after sort #4 from Generation #2 (non-template clones).** Shows convergence on E6R, F19L, and S39P mutations.

**SI References**


1. Turbo
2. Lov turbo
3. G. Chao, W. L. Lau, B. J. Hackel, S. L. Sazinsky, S. M. Lippow, K. D. Wittrup, Isolating and engineering human antibodies using yeast surface display. *Nat. Protocols*, ***1*****(2)**, 755-768, (2006).

4. D. W. Colby, B. A. Kellogg, C. P. Graff, Y. A. Yeung, J. S. Swers, K. D. Wittrup Engineering antibody affinity by yeast surface display. *Methods Enzymol.*, ***388*****(2000)**, 348–358 (2004).
5. Simpson, W. J., and J. R. N. Hammond. *The response of brewing yeast to acid washing*. *Journal of the Institute of Brewing* 95 (1989): 347–54.
6. W. J. Simpson, J. R. M. Hammond, The Response of Brewing Yeasts To Acid Washing. *J. Inst. Brew.*, ***95*****(5**), 347–354 (1989).
7. J. Brown, C. Delaine, O. J. Zacheo, C. Siebold, R. J. Gilbert, G. Van Boxel, A. Denley, J. C. Wallace, A. B. Hassan, B. E. Forbes, E. Y. Jones. Structure and functional analysis of the IGF-II/IGF2R interaction. *EMBO*, **27(1)**, 265–276 (2008).
8. C. Delaine, C. L. Alvino, K. A. McNeil, T. D. Mulhern, L. Gauguin, P. De Meyts, E. Y. Jones, J. Brown, J. C. Wallace, B. E. Forbes. A novel binding site for the human insulin-like growth factor-II (IGF-II)/mannose 6-phosphate receptor on IGF-II. *JBC*, ***282*****(26)**, 18886–18894 (2007).
9. P. C. Fridy, Y. Li, S. Keegan, M. K. Thompson, I. Nudelman, J. F. Scheid, M. Oeffinger, M. C. Nussenzweig, D. Fenyö, B. T. Chait, M. P. Rout, A robust pipeline for rapid production of versatile nanobody repertoires. *Nat. Methods*, ***11*****(12)**, 1253–1260 (2014).
